# Brain-wide topographic coordination of traveling spiral waves

**DOI:** 10.1101/2023.12.07.570517

**Authors:** Zhiwen Ye, Alexander Ladd, Nancy MacKenzie, Ljuvica Kolich, Anna J. Li, Daniel Birman, Matthew S. Bull, Tanya L. Daigle, Bosiljka Tasic, Hongkui Zeng, Nicholas A. Steinmetz

## Abstract

Traveling waves of activity are a prevalent phenomenon within neural networks of diverse brain regions and species, and have been implicated in myriad brain functions including sensory perception, memory, spatial navigation, and motor control. However, their spatial organization, anatomical basis, and whether they are locally confined versus distributed across the brain, remains unclear. Here we used cortex-wide imaging and large-scale electrophysiology in awake mice to reveal the organization of traveling waves across spatial scales. Traveling waves formed spiral patterns predominantly centered on somatosensory cortex, sweeping across somatotopic maps. Strikingly, the local axonal architecture of neurons in sensory cortex exhibited a matching circular arrangement. At the cortex-wide scale, these spiral waves were mirrored between hemispheres and between sensory and motor cortex, reflecting topographic long-range axons. At the brain-wide scale, cortical spiral waves were coordinated with subcortical spiking patterns in the thalamus, striatum, and midbrain. These results establish that traveling waves are shaped by axonal pathways into coordinated spiral patterns that globally impact neural activity across diverse brain systems, revealing a distributed, multi-sensory organizational principle for propagating neural activity.

## Main Text

Spiral waves, also called vortices or rotating waves, are among the dynamic patterns of brain activity (*1–4*) previously observed across various timescales and species including turtles (*5*), mice (*6–8*), rats (*9*), cats (*10*), monkeys (*11*) and humans (*12–14*). Traveling waves, including spirals, are strongly correlated with a variety of perceptual, cognitive, and motor behaviors (*11*, *15–23*). Previous studies of such waves often relied on measures of neural activity at restricted temporal (*13*, *14*) or spatial (*11*, *12*, *15*) resolutions and scales, limiting their ability to identify organizational principles and link these dynamic patterns with underlying features of the neural architecture.

Within the cortex, sensory processing is traditionally viewed as modular and hierarchical, with information processed in parallel sensory systems (visual, auditory, whisker, etc.) and integrated in higher-order association areas (*24*). While evidence suggests multisensory integration occurs at the primary sensory cortical level, such as auditory-visual (*25*, *26*) and visual-somatosensory interactions (*27*, *28*), these few well-studied pathways may constitute only a fraction of the interactions between low-level sensory areas (*28*). Adjacent sensory areas (upper limb, lower limb, mouth, nose, etc) in mammals form a somatotopic homunculus (*29–31*), but the extent to which activity propagates across these regions remains largely unknown.

Considering a broader spatial scale, prior studies of traveling waves in mammals have primarily focused on waves in isolated brain regions, especially neocortex (*5*, *7*, *9*, *11*, *12*, *14*, *15*, *17*, *22*, *23*), hippocampus (*19–21*), and others (*18*, *32*). Despite the known anatomical (*30*, *33–35*) and functional connectivity (*36–38*) between the neocortex and the thalamus, basal ganglia, and midbrain, it remains unclear whether fast traveling waves, on the sub-second timescale of perception and action, are topographically coordinated in relation to long-range connectivity in the adult brain (*8*, *39–43*). The mere existence of axonal projections from one brain region to another does not imply that all patterns of activity are transmitted from one region to the other. Specifically, traveling waves could be in the ‘null space’ of activity, that is, they could be patterns of activity that are not transmitted to target areas (*44*, *45*), for example, if projection neurons do not encode traveling waves. Determining whether traveling waves are indeed localized phenomena that can be adequately studied with a focus on isolated brain regions, or whether understanding their mechanisms and function requires a more distributed perspective, remains unclear.

Here, we used fast mesoscopic imaging with large-scale electrophysiology to elucidate the organization of traveling waves in the awake mouse, the features of axonal architecture underlying them, and their coordination across diverse brain systems.

### Spiral waves in mouse cortex

To investigate the mesoscale spatiotemporal dynamics of neural activity across the cortex, we performed widefield calcium imaging in awake, head-fixed mice expressing either GCaMP7 in excitatory neurons or GCaMP8 in all neurons. We focused our analysis on the 2-8 Hz frequency range, and observed prominent activity in this range across the dorsal cortex during quiet wakefulness states, with the highest power in the retrosplenial cortex (fig. S1). Activity in this frequency range includes an alpha-like 3-6 Hz oscillation in visual cortex (*46–48*) which is correlated with perceptual sensitivity in discrimination tasks (*49*), and may include homologues of the human somatosensory mu rhythm (*50*). To visualize the spatiotemporal propagation of traveling waves in this frequency range, we applied the Hilbert transform to the filtered data for each pixel independently to extract the oscillation phase at each moment in time (Fig. 1A).

**Fig. 1.**
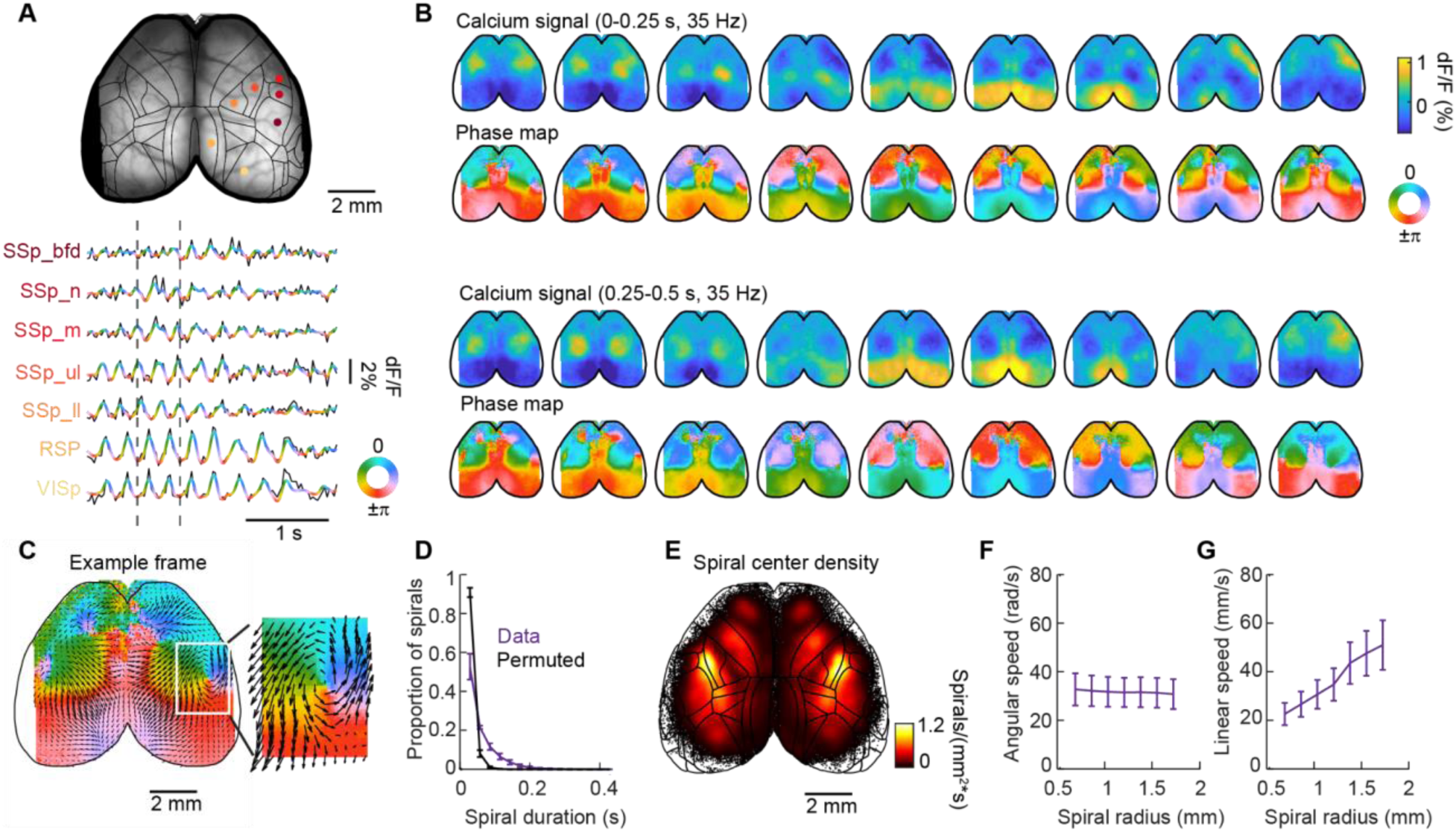
Spiral waves are prevalent in the neural dynamics of mouse cortex. (**A**) Example activity traces from calcium imaging of the dorsal cortical surface. Raw traces (black) are overlaid with 2-8 Hz filtered traces color coded by phase values after the Hilbert transform. (**B**) 18 example consecutive frames (0.5 second) of calcium signal maps and phase maps, within the window indicated by the dashed lines in panel A. (**C**) Phase map from an example frame with spiral in the right hemisphere, and optical flow overlaid depicting wave direction. (**D**) Durations of spirals in data (purple) and in permuted data (black). Error bar: standard error of the mean (n = 15 mice). (**E**) Density estimate of spiral center distribution after automated spiral detection algorithm (n = 15 mice). (**F**) Angular speed distribution as a function of spiral radius across 15 mice. (**G**) Linear speed of spirals as a function of spiral radius. Abbreviations: SSp_bfd (primary somatosensory area, barrel field); SSp_n (primary somatosensory area, nose); SSp_m (primary somatosensory area, mouth); SSp_ll (primary somatosensory area, lower limb); SSp_ul (primary somatosensory area, upper limb); RSP (retrosplenial cortex); VISp (primary visual cortex).

We consistently observed spiral waves, which exhibited a circular arrangement of oscillation phases, in the widefield imaging data (Fig. 1A to C, and fig. S1; Video1). To characterize their properties, we designed an automated spiral detection algorithm (fig. S2). To determine whether observed spirals occurred more frequently than expected by chance, we constructed 3D surrogate data with a Fourier phase shuffling method, in which the spatial and temporal autocorrelations of the data were preserved (Methods). While small spirals commonly occurred by chance in the shuffled data, large spirals (radius >= 40 pixels, or 0.69 mm) occurred in the original data significantly more often than in the shuffled data across sessions (n= 15 mice) and were therefore included for further analysis (fig. S3). Spirals formed sequences persisting for tens to hundreds of milliseconds (Fig. 1B and D). Since single-frame spirals occurred frequently by chance in permuted data (Fig. 1D; Methods), we further focused analyses on spiral sequences lasting multiple sampling frames (>=57 ms; Fig. 1D). Spiral waves with similar characteristics were also observed in LFP activity in wild type mice (fig. S4), indicating that they did not arise from any artifact of calcium imaging. Spirals were also observed in mice that expressed calcium-sensitive fluorescent proteins only in neuron somas and proximal dendritic segments (fig. S5), indicating that they reflected patterns of local activity rather than axonal inputs.

The spiral activity patterns we observed were consistently localized, frequently occurring, and had a characteristic rotational speed. The majority of spiral waves swept across the somatotopic maps of the mouse body representation sequentially, including lower limb, upper limb, mouth, nose and the barrel field (Fig. 1A to C, and fig. S1). Spiral centers were highly concentrated in the middle of primary somatosensory cortex (SSp) across animals (Fig. 1E), a property observed across different mouse lines (fig. S5), spiral sizes, and durations (fig. S6). Spirals had a peak rate of 1.2 spirals/(mm^2^*s) on average, or 3.4% of frames/mm^2^ at 35 Hz (Fig. 1E, and fig. S5, n= 15 mice). Spiral centers often drifted within a spiral sequence and were slightly more likely to rotate counterclockwise in the right hemisphere (fig. S7). Spirals were more prevalent when 2-8 Hz power was high, as were plane waves (fig. S8). However, with improved kinetics of calcium sensors in GCaMP8m compared to GCaMP7, occurrence of detected spiral waves increased (GCaMP8m: 10 ± 0.9%; GCaMP7: 2 ± 0.2%) while plane waves decreased (GCaMP8m: 13 ± 0.9%; GCaMP7: 23 ± 2%; fig. S5 and fig. S8). The occurrence of spiral waves were similarly prevalent when subjects were active or quiescent (fig. S8). Spiral waves had consistent angular speed (*ω)* across different radii (31.9 ± 6.1 rad/s, mean ± STD, equivalent to 5.1 ± 1.0 Hz; Fig. 1F, and fig. S9) but increasing linear speed as a function of radius (Fig. 1G, and fig. S9). The propagation speeds of these waves (0.05 ± 0.01 m/s at 1.7 mm spiral radius, mean ± STD) are therefore substantially slower than axonal conduction velocities, which are on the order of 0.5 m/s for lateral axons in rodent neocortex (*51*), and likely reflect axon conduction across multiple connections.

### An axonal basis for spirals in sensory cortex

To investigate the anatomical basis for the propensity of spirals to form around the somatosensory cortex, we examined the local axonal architecture of neurons in sensory cortex. Using an open dataset of cortical neuron reconstructions (*52*), we discovered that local axons from mouse cortical neurons are also organized circularly (Fig. 2A to C). In particular, the major axes of the axonal terminals from single neurons exhibited a strong tendency to align perpendicular to the vector from the SSp center to the soma location (Fig. 2C and E; 87.4°, n = 435 neurons; 95% confidence interval for circular t-test = 82.8°-92°, *p* < 0.05; real data vs permutation distribution: *p* < 0.05, Watson’s U^2^ test). Moreover, the orientation of these axonal axes matched the propagation directions of spiral waves, exceeding a permuted distribution of flow vectors (Fig. 2D and F; see Methods; *p* = 0.004, student’s t-test). We did not observe a circular arrangement of axons in motor cortex, though there was a non-random organization with a bias in the anterior-posterior direction (fig. S10). A computational simulation of activity propagation over sheets of cortical neurons, modeled as coupled oscillators with circularly biased connectivity as in the data, exhibited strengthened spiral dynamics relative to a model with isotropic connectivity (fig. S10). The anatomical architecture of axons in the somatosensory cortex therefore corresponds to the spiral waves we observed, and could support the occurrence of these dynamic patterns.

**Fig. 2.**
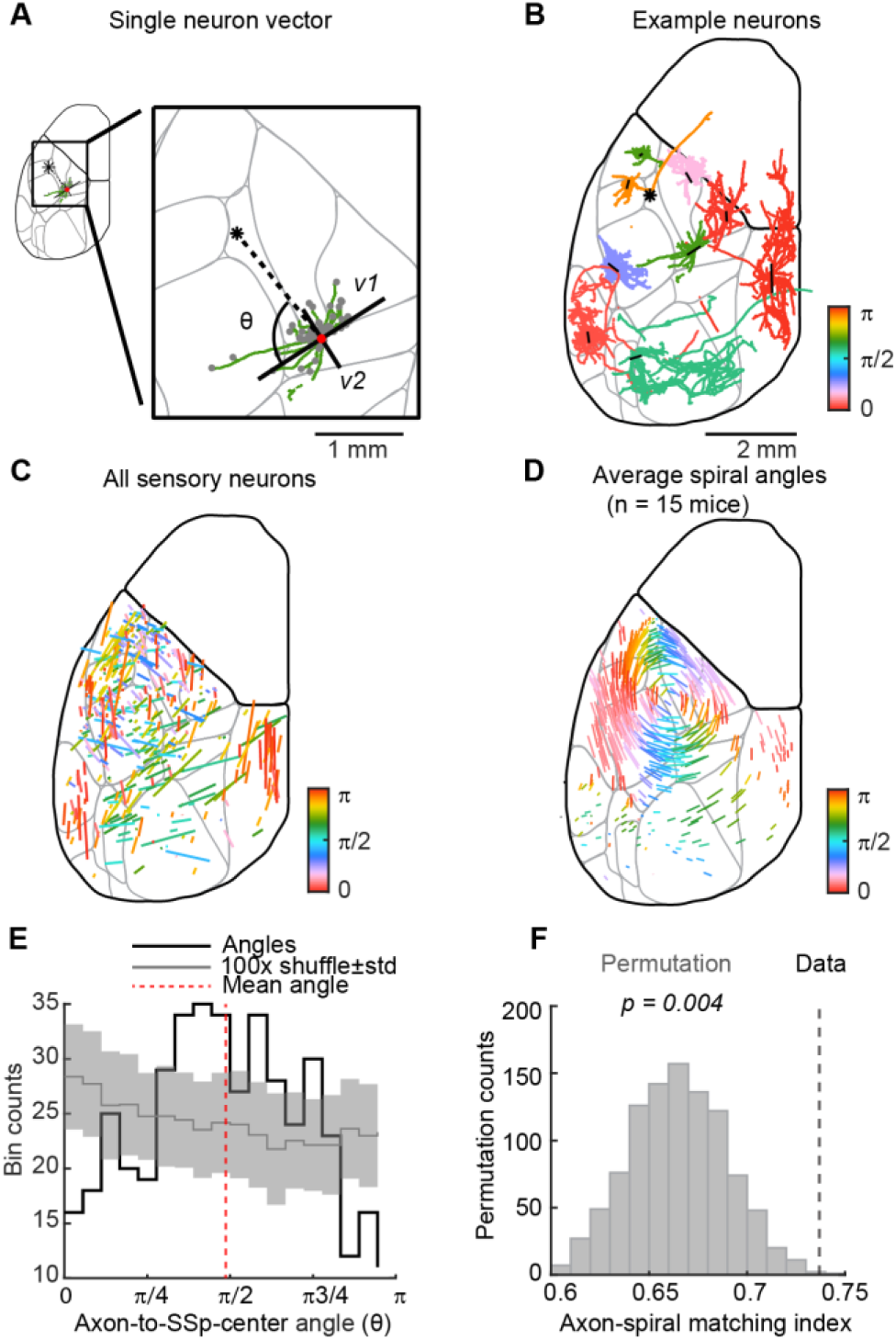
Axon orientations of single neurons match the circular arrangement of spiral waves around the middle of somatosensory cortex. (**A**) Morphology of an example neuron’s axonal arbor (green) with local axon terminals overlaid (gray). The top two singular vectors of the terminal point cloud (*v1*, *v2*; black) are overlaid on top of the axonal arbor and centered on the soma (red). Asterisk represents the middle of SSp, within the unassigned SSp (SSp-un) zone. Axon-to-SSp-center angle (θ) is the angle between the first singular vector (*v1*) and the vector from the soma to the SSp middle. (**B**) Axonal arbors of 9 example neurons. The direction of the vector overlaid on top of the arbor (black) represents the first singular vector of the axon terminal cloud. The length of the vector represents axonal arbor polarity, which is the ratio between the first two singular values, with longer length indicating higher polarity. The vectors are centered on the cell soma positions. (**C**) Axon orientation and polarity for all sensory neurons (n = 435 neurons), as in B. Color represents the angle of the vectors. (**D**) Mean activity flow vectors of a collection of large spirals with the same direction of rotation across mice (n = 15). (**E**) Histogram of axon-to-SSp-center angle (θ) for all 435 neurons. The mean angle across all neurons is indicated by the dashed red line. Mean and standard deviation of shuffled distribution is indicated in gray. (**F**) Axon-flow matching index is significantly greater than the permuted distribution of flow vectors.

### Topographically mirrored spirals across cortex

Spirals were often symmetrical between left and right hemispheres, as well as between sensory and motor cortex, divided along the border of primary somatosensory cortex (SSp) and primary motor cortex (MOp). When 2 spirals were detected simultaneously in neighboring mirrored brain structures at the same time (SSp-MOp, SSp-left-right, MOp-left-right), spirals were mostly traveling in the opposite direction (Fig. 3A; full comparison, fig. S7).

**Fig. 3.**
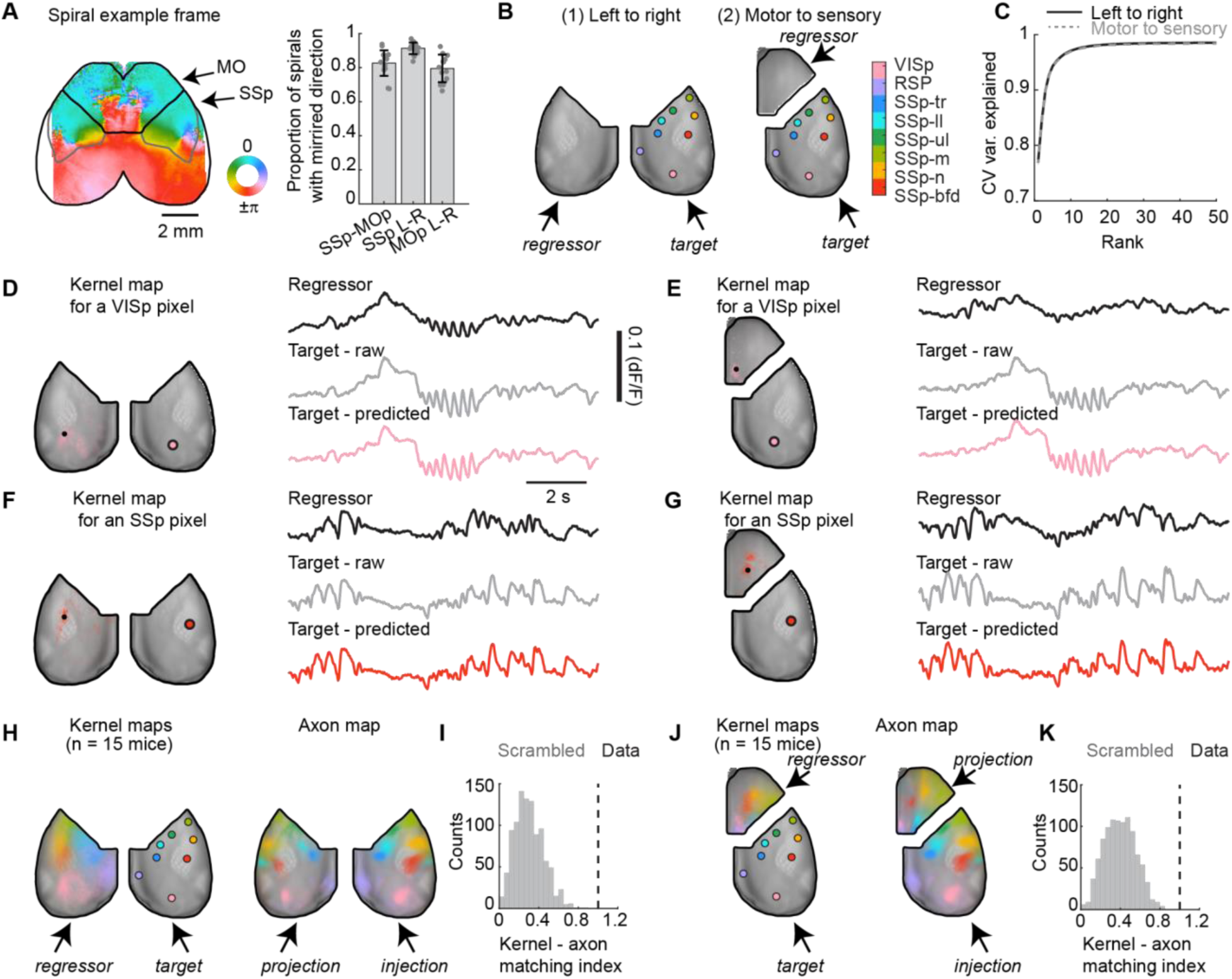
Spirals are mirrored between left and right hemispheres, and between sensory and motor cortex, reflecting an underlying topographic mapping of activity and long-range axons. (**A**) Left: an example phase map with SSp/MOp border depicted in black outline. Right: proportion of paired spirals with mirrored direction between topographically connected cortical regions, including SSp versus MOp on the same hemisphere, SSp on left versus right hemispheres (SSp L-R), and MOp on left versus right hemisphere (MOp L-R). (**B**) Schematic of regressions to predict right sensory cortical activity from (1) left sensory cortex and (2) right hemisphere motor cortex. Centers of 8 sensory cortical regions are highlighted in colored dots. (**C**) Variance explained plateaued with around 16 latent components with Reduced Rank Regression, for both left-to-right prediction and motor-to-sensory prediction. (**D**) Left: regression kernel map in the left hemisphere for predicting an example right VISp pixel. The “kernel map” is a visualization of the predictor weights with each pixel’s color transparency scaled by the value of the weight matrix. Right: example regressor trace (black) with the highest value in the weight matrix from left hemisphere (top right); raw (gray, middle right) and predicted activity trace (pink, bottom right) of the example right VISp pixel. (**E**) Same as D, but with motor cortex traces as regressors. (**F**) Same as D, but for predicting an SSp-bfd pixel. Example trace was from the same time epoch as D, E. (**G**) Same as E, but for predicting an SSp-bfd pixel with motor cortex traces as regressors. (**H**) Left: averaged kernel maps for all example pixels in the right sensory cortex across 15 mice. Right: Axon targets in the left hemisphere from viral injections in the right hemisphere (*35*). (**I**) Topographic kernel-axon matching index, measuring how similar the axon projection fields are to the kernel maps in H, is significantly greater than the distribution of the index when permuting the injection identities. (**J**) Same as H, but for activity kernels of motor-to-sensory prediction and axon targets in the motor cortex from the same viral injections as in H. (**K**) Same as I, but for kernel maps and axon project fields in J.

With the observation of cortical phase map symmetry along the border of SSp and MOp (Fig. 1B, Fig. 3A), we sought to look for the basis of spiral symmetry with functional connectivity. We used a reduced rank regression model to (1) predict activity in the right hemisphere from the left hemisphere or (2) predict activity in the sensory cortex from the motor cortex, divided along the border of SSp and MOp (Fig. 3B). Reduced-rank regression analysis achieved 98 ± 0.1% (mean ± SEM, 15 mice) cross-validated variance explained with 16 latent reduced-rank components for both left-to-right and motor-to-sensory predictions (Fig. 3C). As a result, neural activity of a given pixel in one hemisphere (or sensory cortex) can be fully represented as a linear combination of neural activity in the other hemisphere (or motor cortex) with its own unique and spatially localized kernel map, i.e., weight matrix (Fig. 3D to G). Therefore, moment-to-moment population activity information is topographically shared between hemispheres, as well as between the sensory and motor cortex.

The spatial arrangement of the localized kernel maps closely matched the axonal projection maps across hemispheres (Fig. 3H, and fig. S11), significantly exceeding scrambled axonal distribution (Fig. 3I; kernel-axon matching index for real data: 1; 1000x permutation index: 0.3 ± 0.005, mean ± SEM; *p* = 1.6 x 10^-6^; Student’s t-test; Methods). The same matching patterns were observed between activity kernel maps and axon projection maps between sensory and motor cortex (Fig. 3J and K, and fig. S11; kernel-axon matching index for real data: 1; 1000x permutation index: 0.4 ± 0.005, mean ± SEM; *p* = 1.4 x 10^-4^; Student’s t-test).

### Brain-wide coordination of spiral waves

To investigate whether individual spiral waves simultaneously occur in cortex and in subcortical areas, we combined widefield imaging in the cortex with 4-shank Neuropixels electrophysiology in subcortical areas, including thalamus, striatum and midbrain (Fig. 4A). When 2-8 Hz oscillation was prominent in the cortex, we consistently observed corresponding oscillations in the subcortical spiking activity (Fig. 4B and C, and fig. S12). To test whether subcortical spiking activity contains information about cortical spiral wave propagation, we implemented a linear regression model to predict cortical activity from subcortical spiking activity. Subcortical spiking could predict cortical activity well across much, but not all, of the cortical surface (Fig. 4B and C, and fig. S13), presumably because the subcortical regions were incompletely sampled.

**Fig. 4.**
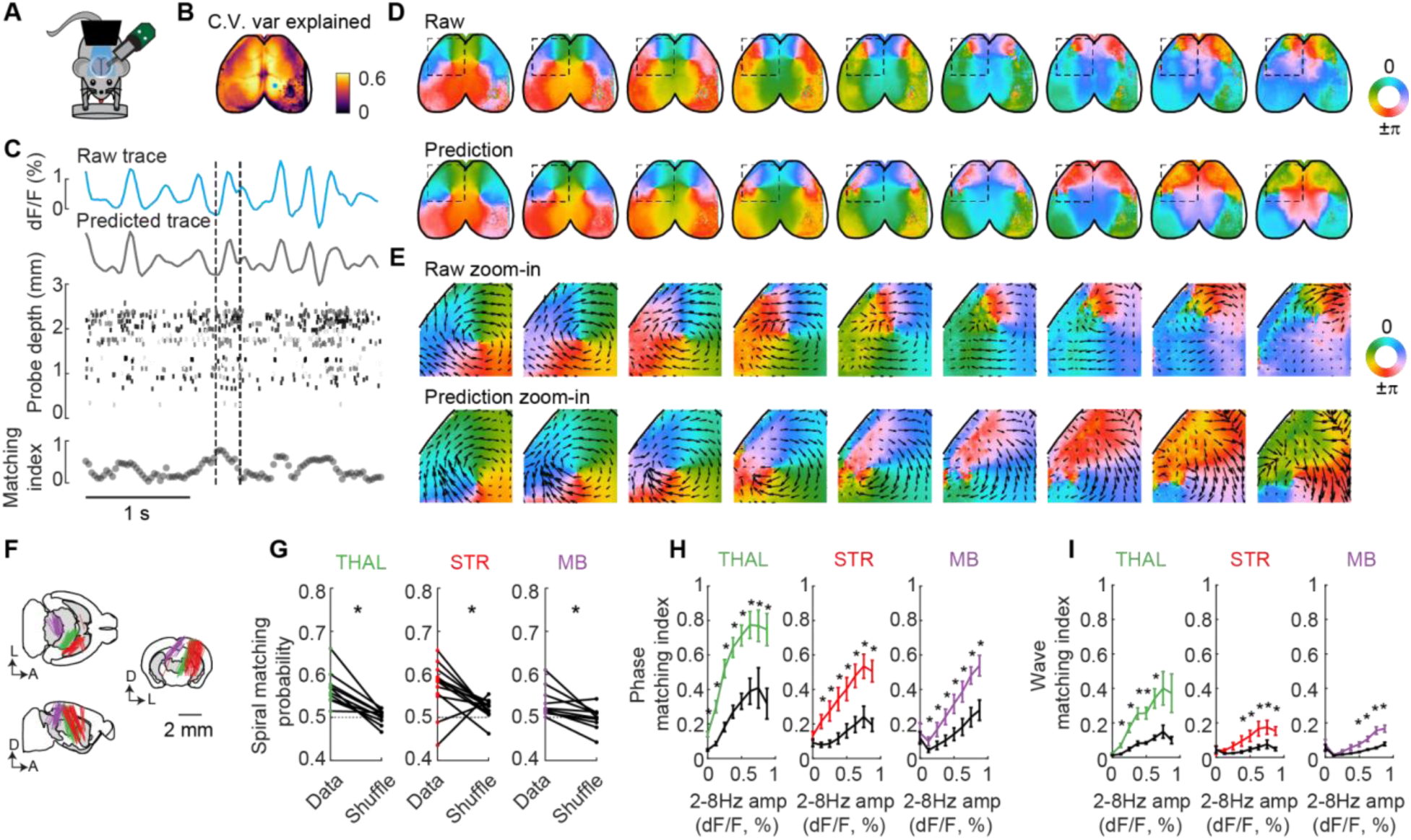
Spirals are coordinated between cortical and subcortical regions. (**A**) Illustration of simultaneous cortical widefield and subcortical electrophysiological recordings with 4-shank Neuropixels 2.0 probes. (**B**) Example map showing cross-validated variance explained of cortical widefield data predicted from thalamic population spiking. The craniotomy was in the right posterior cortex. (**C**) Top: an example epoch of raw (blue) and predicted (gray) cortical calcium activity for an example pixel highlighted in B. Middle: thalamic spike raster plot, ordered by probe depth. The tips of the 4-shank probe were at 0 mm. Each neuron was color coded by a randomized shade of gray color. Bottom: Traveling wave matching index, comparing optical flow vectors between raw and predicted phase maps. (**D**) Example raw (top) and predicted (bottom) phase maps of cortical activity within the dashed lines in C. (**E**) Zoom-in of raw (top) and predicted (bottom) phase maps within the square ROI in D, with activity flow vectors overlaid. (**F**) Probe trajectories were retrieved successfully in 30/32 sessions (thalamus: 10, striatum: 12, midbrain 8/10). (**G**) Spiral matching probabilities between spirals detected from raw and predicted cortical phase maps were above chance level (0.5), and were significantly greater than spiral matching probabilities from predictions with shuffled neuron identity across sessions. (**H**) Predicted phase maps were more likely to match raw phase maps as 2-8 Hz amplitude increased. Phase matching indexes from subcortical spiking prediction were significantly higher than predictions from shuffled neuron identities. (**I**) Direction of predicted traveling waves were more likely to match raw traveling waves as 2-8 Hz amplitude increased. Wave matching indexes were significantly higher than predictions from shuffled neuron identities.

We assessed whether spirals were present in the activity patterns of subcortical areas by asking whether predictions from subcortical activity recapitulated spirals in the cortical activity (Fig. 4D and E, and fig. S12). We computed the probability that spirals in the raw cortical frames rotated in the same direction as spirals in the predicted frames from all thalamic, stiatal and midbrain spiking data (Fig. 4F). Spirals were significantly more likely to match rotation directions from prediction with the real spiking data compared to data in which neuron identities were shuffled, and the matching probability was above chance level (thalamus: 0.57 ± 0.01 vs 0.5 ± 0.01, *p* = 3.8×10^-4^; striatum: 0.57 ± 0.02 vs 0.52 ± 0.01, *p* = 0.03; midbrain: 0.53 ± 0.01 vs 0.5 ± 0.01, *p* = 0.008; mean ± SEM; paired Student’s t-test) for all subcortical regions (Fig. 4G).

To determine whether traveling waves were more generally coordinated outside of spirals, we quantified waves with optical flow and compared them between raw and predicted cortical activity (Fig. 4D and E, and fig. S12). Firstly, as cortical 2-8 Hz oscillation amplitude increased, phase maps were more likely to match for raw cortical activity and predicted activity from all subcortical regions across sessions (Fig. 4H; thalamus: *p* = 6×10^-20^; striatum: *p* = 1×10^-6^; midbrain: *p* = 2×10^-12^; main effect of 2-8 Hz amplitude, unbalanced two-way ANOVA). Moreover, as cortical 2-8 Hz oscillation amplitude increased, traveling wave directions were also more likely to match for cortical activity and predicted activity from all subcortical regions (Fig. 4I; thalamus: *p* = 3×10^-15^; striatum: *p* = 5×10^-4^; midbrain: *p* = 8×10^-15^; main effect of 2-8 Hz amplitude, unbalanced two-way ANOVA). Probability of wave matching significantly exceeded shuffled permutation for the majority of single sessions (fig. S13; 10/10 sessions in thalamus, 12/12 sessions in stratum and 9/10 sessions in midbrain; unbalanced two-way ANOVA) and across sessions (Fig. 4I; thalamus: *p* = 1×10^-16^; striatum: *p* = 2×10^-6^; midbrain: *p* = 7×10^-10^; unbalanced two-way ANOVA). Therefore, cortical spirals and traveling waves are represented in subcortical brain areas on a moment-to-moment basis.

## Discussion

We observed frequently occurring spiral waves traveling across the somatotopic maps of the mouse sensorimotor cortex, matching a circular pattern of local axonal architecture. Spirals were also topographically mirrored between left and right cortical hemispheres, and between sensory and motor cortex, respecting long-range axonal connectivity. Spirals were widely distributed and coordinated across cortex, striatum, thalamus, and midbrain. These results establish that traveling waves in the mammalian brain are organized as large-scale spiral waves. Moreover, they are not merely a local phenomenon but are globally coordinated across the brain, and therefore cannot be adequately understood by studying any one brain region in isolation.

Beyond the established modular and hierarchical sensory processing pathways (*24*, *28*), we provide evidence that traveling spiral waves propagate across the sensory cortices with an underlying circular axonal connectivity, potentially facilitating cross-module interactions at the primary sensory level. Spiral waves were prevalent (10 ± 0.9% of frames in GCaMP8m mice; fig. S5 and fig. S8) and propagated across somatosensory modalities, including the lower limb, upper limb, mouth, nose and barrel field. Visual and auditory cortices also integrate into the larger spiral pattern, encompassing the entire sensory cortex (Fig. 2). Previously reported cross-modal interactions (*25–28*) may therefore reflect specific subsets of these broader wave-based interactions that sweep across multiple modalities. At the neuronal level, individual neurons vary in their coupling to population activity (*53*), but the identity of neurons that are modulated by traveling waves, the nature of information borne on the waves, and whether neuronal participation dynamically changes with behavior and learning remain open questions.

We found that spirals were centered in the middle of somatosensory cortex, an area of “unassigned” function in the Allen CCF (”SSp-un”) (*55*) and unique in its poorly defined layer 4 compared to other somatosensory regions (*56*, *57*). This region may therefore comprise a discontinuity or ‘defect’ in the cortical medium. Our findings show that cortical spirals are closely linked to local axonal architecture. However, the direction of causality between anatomy and function remains to be tested, that is, whether the presence of circularly arranged axonal architecture drives spiral activity, or whether spiral activity patterns in this region drive the formation of this axonal architecture via plasticity mechanisms during development (*42*, *58*).

The observation of matching waves in three subcortical regions raises the possibility that the medium of wave propagation may not be limited to corticocortical connectivity. For example, thalamic mechanisms may contribute to global wave propagation (*47*, *59–63*). In particular, lateral connectivity within the reticular nucleus of thalamus or connectivity linking cortical regions via thalamic intermediaries might play a role coordinating recurrent thalamocortical activity (*64–68*). Future studies of the mechanisms of traveling waves in the mammalian brain will need to consider the diverse circuits across the brain that simultaneously reflect their propagation. Whether dynamics beyond the frequency ranges we studied here on faster (*11*, *15*, *69*) or slower (*13*, *14*) timescales follow the same principles is an open question, as some evidence suggests that cortical activity in disparate frequency ranges may be uncoupled (*70*). Moreover, the presence of these waves in basal ganglia and midbrain, regions traditionally more tightly linked with motor function, raises the question of how these activity patterns avoid interfering with motor control, and future studies may seek to establish with the methods employed here whether these traveling waves are indeed even present at the level of pons, hindbrain, or spinal cord (*71*).

What functions might these spiral dynamics serve? Spiral waves may act as spatiotemporal clocks, coordinating neuronal spiking to encode sequential sensation and action of the internal body, akin to hippocampal waves representing external space or events (*19*). Second, these waves could be involved in guiding plasticity mechanisms (*2*) to form connections between neighboring somatosensory and motor modalities (e.g., hindlimbs with forelimbs, trunk area with facial area, etc.) to support behavioral functions that commonly involve sequential sensory and motor behaviors across these body parts (*22*, *23*, *72*). The spirals we observed also have similar properties to those observed during human sleep spindles, suggesting a possible role in memory consolidation (*12*). Third, oscillatory power in the frequency range that we studied is correlated with behavioral engagement (*49*), and the momentary phase of spontaneous fluctuations modulates perceptual detectability (*15*, *16*). Therefore, we hypothesize that waves of activity sequentially sweeping across sensory modalities may act to continuously scan these sensory systems, like a radar sweep (*1*, *73*), modulating stimulus detectability in each region while maintaining metabolic efficiency. Considered broadly, topographically coordinated spiral waves across cortical and subcortical regions may provide the scaffold for a diverse array of perceptual and cognitive functions.

## Acknowledgements

We thank Sverre Grødem, Kristian Lensjø, and Marianne Fyhn for the GCaMP8 viruses. We thank Sam Golden for lightsheet imaging support. We thank Bing Brunton, Kenneth Harris, Ryan Raut, Harsha Gurnani, and members of the Steinmetz Lab for discussions about this work.

## Author contributions

ZY and NAS conceived and designed the study.

ZY performed experiments and data analysis.

MSB, ZY, AL and NM conducted modeling work.

LK assisted animal husbandry.

TLD and BT created Ai195 and Ai210 transgenic mouse lines.

AJL assisted facial motion energy analysis.

AJL and DB helped develop experimental protocols.

ZY and NAS wrote the original draft manuscript.

ZY, MSB, AL, DB and NAS revised the manuscript.

HZ obtained funding for creating transgenic mouse lines.

NAS obtained primary funding for the project.

## Data and materials availability

All data are available at: https://doi.org/10.6084/m9.figshare.27850707.v1. A detailed data description and companion code are at: https://github.com/zhiwen10/YE-et-al-2023-spirals.

## Materials and Methods

### Animals and virus injections

All experimental protocols were conducted according to US National Institutes of Health guidelines for animal research and approved by the Institutional Animal Care and Use Committee at the University of Washington.

The following transgenic mouse lines and viruses were used for this study (Table 1). Mouse lines:

**Table 1.** Transgenic mice and viral injection strategies.

| Mouse ID | Sex | Age (days) | GCaMP | Genotype | Virus injection | WF sessions | WF+ephys sessions |
| --- | --- | --- | --- | --- | --- | --- | --- |
| AB_0004 | M | 65 | jGCaMP7f | Ai210 | 5μL PHP.eB-flpO | 1 | 0 |
| ZYE_0012 | F | 50 | jGCaMP7f | Ai210 | 5μL PHP.eB-flpO | 1 | 0 |
| ZYE_0043 | F | 104 | jGCaMP7f | Ai210 | 5μL PHP.eB-cre | 1 | 1 |
| ZYE_0052 | M | 86 | jGCaMP7f | Ai210 | 5μL PHP.eB-cre | 1 | 1 |
| ZYE_0053 | M | 88 | jGCaMP7f | Ai210 | 5μL PHP.eB-cre | 0 | 3 |
| ZYE_0056 | F | 109 | jGCaMP7f | Ai210 | 5μL PHP.eB-cre | 1 | 0 |
| ZYE_0058 | M | 167 | jGCaMP7f | Ai210 | 5μL PHP.eB-cre | 1 | 2 |
| ZYE_0059 | M | 134 | jGCaMP7f | Ai210 | 10μL PHP.eB-cre | 1 | 2 |
| ZYE_0060 | M | 143 | jGCaMP7f | Ai210 | 10μL PHP.eB-cre | 1 | 2 |
| ZYE_0062 | F | 149 | jGCaMP7f | Ai210 | 10μL PHP.eB-cre | 1 | 3 |
| LK_0003 | M | 72 | jGCaMP7f | Ai210 triple | / | 1 | 1 |
| ZYE_0048 | F | 154 | jGCaMP7f | Ai210 triple | / | 0 | 2 |
| ZYE_0050 | F | 148 | jGCaMP7f | Ai210 triple | / | 0 | 2 |
| ZYE_0014 | F | 235 | jGCaMP7s | Ai195 | 3μL PHP.eB-flpO | 0 | 1 |
| ZYE_0038 | F | 79 | jGCaMP7s | Ai195 | 5μL PHP.eB-cre | 0 | 1 |
| ZYE_0041 | F | 122 | jGCaMP7s | Ai195 | 5μL PHP.eB-cre | 0 | 4 |
| ZYE_0047 | M | 74 | jGCaMP7s | Ai195 | 5μL PHP.eB-cre | 0 | 1 |
| LK_0001 | M | 112 | jGCaMP7s | vglut1-cre | 40μL PHP.eB- flex-g7s | 1 | 0 |
| ZYE_0066 | F | 236 | jGCaMP8m | wild type | 100μL CAP-B10-G8m | 0 | 1 |
| ZYE_0067 | F | 234 | jGCaMP8m | wild type | 100μL CAP-B10-G8m | 1 | 1 |
| ZYE_0070 | F | 102 | jGCaMP8m | wild type | 100μL CAP-B10-G8m | 1 | 0 |
| ZYE_0071 | F | 101 | jGCaMP8m | wild type | 100μL CAP-B10-G8m | 1 | 1 |
| ZYE_0072 | M | 102 | jGCaMP8m | wild type | 100μL CAP-B10-G8m | 1 | 0 |
| ZYE_0087 | F | 269 | GCaMP8s | GCaMP8s | / | 0 | 3 |
| ZYE_0088 | F | 230 | GCaMP8s | GCaMP8s | / | 0 | 3 |

_Ai210 (B6.Cg-*Igs7*_*^tm210(tetO-GCaMP7f,CAG-tTA2)Tasic^*/J, Jax# 037378)

- ● dual-recombinase responsive (cre/flp-dependent), express tTA2 and jGCaMP7f

_Ai195 (B6;129S6-*Igs7*_*^tm195(tetO-GCaMP7s,CAG-tTA2)Tasic^*/J, Jax# 034112)

- ● dual-recombinase responsive (cre/flp-dependent), express tTA2 and jGCaMP7s

tetO-GCaMP8s (B6;D2-Tg(tetO-GCaMP8s)1Genie/J, Jax# 037717)

- ● Transgenic mice expressing jGCaMP8s, controlled by a tetracycline-responsive promoter element (TRE; tetO)

Camk2a-tTA (B6.Cg-Tg(Camk2a-tTA)1Mmay/DboJ, Jax# 007004)

- ● allow the inducible expression of genes in forebrain neurons, when mated to strain carrying a gene of interest under the regulatory control of tetO

Snap25-Cre (B6;129S-*Snap25^tm2.1(cre)Hze^*/J, Jax# 023525)

- ● knock-in mice with widespread Cre recombinase expression directed throughout the brain

VGlut1-FlpO (B6;129S6-*Slc17a7^em1(flpo)Tasic^*/J, Jax# 034422)

- ● knock-in mice with optimized FLP recombinase expression directed to VGlut1-expressing cells

VGlut1-Cre (B6;129S-*Slc17a7^tm1.1(cre)Hze^*/J, Jax# 023527)

Viruses:

- ● knock-in mice with Cre recombinase expression directed to VGlut1-expressing cells

AAV-PHP.eB-GcaMP7s (addgene# 104487-PHPeB): AAV PHP.eB particles produced from pGP-AAV-syn-jGCaMP7s-WPRE (addgene #104487) at titer ≥ 1×10^13^ vg/mL

CAP-B10-soma-jGCaMP8m (gift from Sverre Grødem (*74*)) : AAV with CNS- and neuronal-specific CAP-B10 serotype produced from pGP-AAV-syn-soma-jGCaMP8m-WPRE (addgene #169257) at titer 3.5 x 10^12^ vg/mL

AAV-PHP.eB-hSyn-Cre (addgene # 105540-PHPeB): AAV PHP.eB particles produced from pENN.AAV.hSyn.HI.eGFP-Cre.WPRE.SV40 (addgene #105540) at titer ≥ 1×10^13^ vg/mL

AAV-PHP.eB-Syn-flpO-WPRE (gift from Tanya Daigle, Allen Institute for Brain Science): AAV PHP.eB particles produced from Syn-FlpO-WPRE at titer 4.1×10^13^ vg/mL

We used five different strategies to express GCaMP7/8 across the brain. (1) Ai210 or Ai195 mice were bred with Snap25-Cre or VGlut1-FlpO, then injected with AAV-PHP.eB-Syn-FlpO or AAV-PHP.eB-hSyn-Cre virus (3-5 μL) retro-orbitally in 4-6 week-old mice for dual cre/FlpO-dependent GCaMP7f or GCaMP7s expression throughout the brain; (2) Ai210 mice were bred with Snap25-Cre positive and VGlut1-FlpO positive mice, to create triple transgenic mice that expressed GCaMP7f in excitatory neurons throughout the brain; (3) Retro-orbital injection of CAP-B10-soma-jGCaMP8m (100 μL) in 4-6 week-old wild type mice; (4) Retro-orbital injection of AAV-PHP.eB-flex-GCaMP7s (40 μL) in 4-6 week-old VGlut1-Cre mice; (5) tetO-GCaMP8s mice were bred with Camk2a-tTA mice.

In total, 25 mice of both sexes with GCaMP expression were used, between 2-8 months old (median at 4 months old) when the first experiment was conducted. 15 mice were used for widefield imaging only experiments, 19 mice (32 sessions) were used for simultaneous widefield and electrophysiological recordings. 9 out of the total 25 mice were used for both experiments. Our findings were consistent across different expression strategies, and therefore we combined these mice in the analysis unless specified.

### Surgery

The surgical preparation was similar to that used previously for whole-cortex widefield imaging (*75*). Mice were anesthetized under 3-4% isoflurane in an induction chamber first, then maintained at 1-2% isoflurane for the duration of the 1-2 hour procedure. Carprofen (5 mg/kg) was administered subcutaneously and lidocaine (2 mg/kg) was injected under the scalp for postoperative analgesia. The scalp was shaved and further cleaned with hair removal cream before mice were transferred to a separate stereotaxic frame for surgery. Eye ointment (Alcon, Systane) was applied before surgery to prevent drying and body temperature was kept at 37 °C with a far infrared warming pad (Kent Scientific). The skin and periosteum connective tissue was cleared off the dorsal surface of the skull. A layer of cyanoacrylate (Vetbond Tissue Adhesive) was then applied to seal the junction between the exposed skull and cut skin. A 3D-printed plastic recording chamber was implanted on top of the skull with dental cement (C&B Metabond) to provide light isolation during experiments, and a custom-made steel headplate was attached to the skull over the interparietal bone with Metabond for head fixation. To create a clear coating suitable for imaging, we consecutively applied two separate thin layers of fast curing optical adhesive (NOA81, Thorlabs) and cured until solidification with a 365 nm, 3 W output power UV Flashlight (LIGHTFE, UV301D). Carprofen (0.05 mg/ml) was given for 3 days in water after surgery. Mice were allowed to recover in the home cage for at least 5 days before habituation to head fixation.

### Widefield imaging and data processing

The widefield imaging setup was custom-built with a macroscope body (sciMedia fluorescent beam splitter DLFLSP2R) fitted with a CMOS camera with 3.45 μm/pixel resolution (Basler ace acA2440-75 m) and a 0.63× objective lens (Leica Planapo 0.63×). Alternating dual light illumination was generated using an OptoLED system (Cairn Research) with blue (470 nm) and violet (405 nm) LEDs to capture GCaMP calcium signal and hemodynamic intrinsic signal at 35 Hz each, respectively. Images were captured at 70 Hz using external triggers synchronized to the light illumination and binned at 3 x 3 pixels to increase signal-to-noise ratio and reduce data size. This results in a final resolution of 17.3 μm/pixel for the collected imaging data. To reduce light artifacts for Neuropixels recordings during imaging, the LED onset and offset were modified to be a 3-ms sine-wave ramp by controlling the analog voltage output from a microcontroller board (Teensy 3.2), resulting in an effective 11-ms exposure time at full intensity for each frame.

For widefield imaging experiments, mice were acclimated to head-fixation for at least two sessions before recording, following recovery from headplate implant surgery. Head-fixed mice were seated on a plastic tube with forepaws on a left-right rotating rubber wheel (*76*). Three iPad screens were positioned around the mouse at right angles. A 3D-printed cone was designed to connect the recording chamber with the macroscope objective in order to shield the subject’s eyes from the LEDs and prevent light contamination of the imaging signal. Eye position and facial movements were tracked simultaneously with separate infrared cameras (eye position: Navitar Zoom 7000; facial movement: FLIR Chameleon3) with external infrared illumination (CMVision IR30, 850 nm). Cortical visual areas were mapped using white visual sparse noise squares asynchronously on a black background and visual field sign maps (fig. S5) were computed as in Peters et al. (2021)(*37*).

For widefield data analysis, non-neuronal signals arising from changes in blood flow were removed by subtracting the violet-light evoked signals from the blue-light evoked signals with linear regression, which resulted in an effective sampling rate of GCaMP activity at 35 Hz (*75*, *77*, *78*). In brief, a scaling factor was first fit for each pixel between two signals that were band-pass filtered in the range of 10-13 Hz (heartbeat frequency, with largest haemodynamic effect expected). The haemodynamic-corrected traces were therefore the blue traces in full frequency range subtracted by the scaled violet traces (*37*, *75*). All widefield data were aligned to the Allen CCF (*55*) with affine transformation using four anatomical landmarks as in Musall et al. (2019) (*78*), which were the left, center and right points where the anterior cortex meets the olfactory bulbs, and the medial point at the base of retrosplenial cortex. Alignment results were further verified with retinotopic visual field sign maps, with visual fields aligned within the boundaries of the visual CCF coordinates (fig. S5).

To store and process the data effectively, we compressed the widefield data (D) into spatial components (U) and temporal components (S*V) with singular value decomposition in the form D = USV^T^, as previously described (*37*, *49*). We used the top 50 singular vectors, which accounted for 97.3 ± 0.5% (mean ± SEM, 15 mice) of total variance before compression. We divided the widefield signal by the average fluorescence at each pixel, to get ΔF/F normalized signal.

### Simultaneous Neuropixels recording and widefield imaging

Penetration angles and positions of 4-shank Neuropixels 2.0 probes (*79*) for subcortical recordings were carefully designed to maximize the coverage of sensory modalities in each subcortical region (thalamus, striatum and midbrain) based on the 3D axonal projection maps from the Allen Mouse Brain Connectivity Atlas (*35*), using the neuropixels trajectory explorer (https://github.com/petersaj/neuropixels_trajectory_explorer) and Pinpoint software (*80*). In detail, eight injection experiments with anterograde virus injection sites at VISp [ID: 309003780], RSP [ID: 166054929], SSp-tr [ID: 100141495], SSp-ul [ID: 286312782], SSp-ll [272698650], SSp-m [ID: 157654817], SSp-n [ID: 168163498], SSp-bfd [ID: 127866392] were selected and 3D projection density volumes were downloaded from the Allen Mouse Brain Connectivity Atlas. Intensity-weighted centers of the eight projections within each subcortical region (thalamus, striatum and midbrain) were identified, and a 2D plane was fitted from these center points. Optimal angles and depths of the 4-shank probe were then selected within the fitted 2D plane, which covered most of the projected center points (Table 2).

**Table 2.**
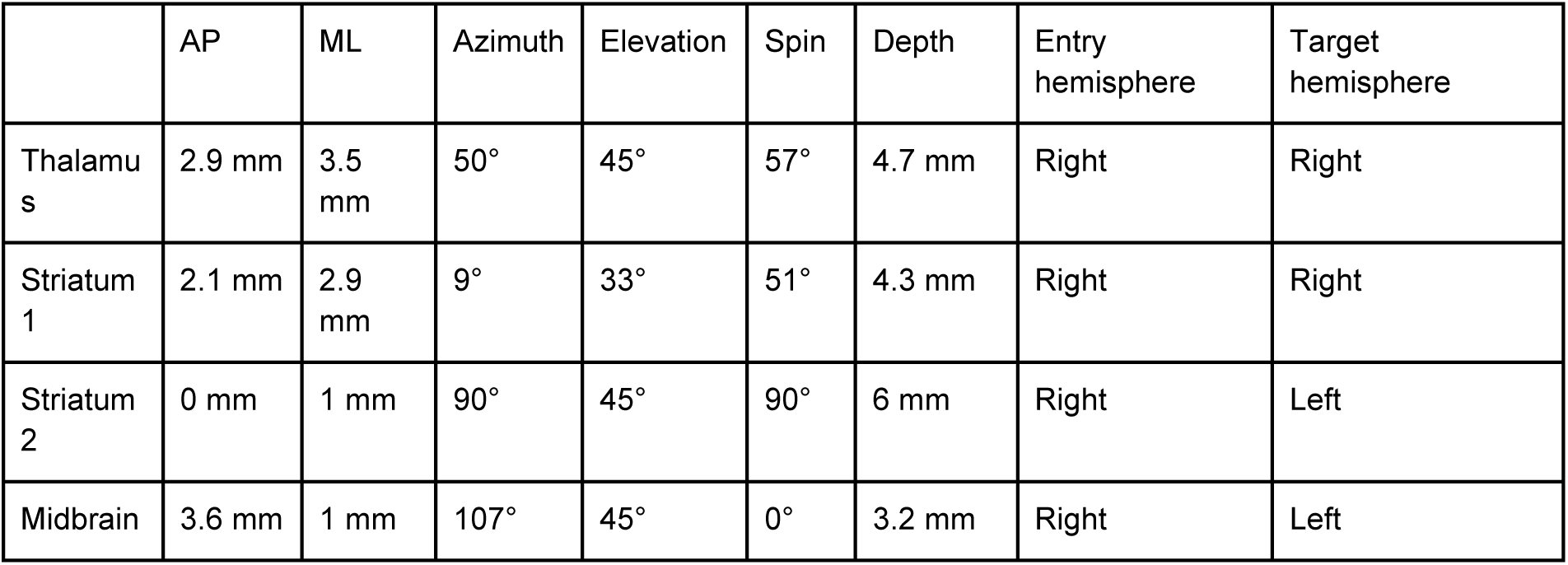
Neuropixels targeting angles.

|  | AP | ML | Azimuth | Elevation | Spin | Depth | Entry hemisphere | Target hemisphere |
| --- | --- | --- | --- | --- | --- | --- | --- | --- |
| Thalamus | 2.9 mm | 3.5 mm | 50° | 45° | 57° | 4.7 mm | Right | Right |
| Striatum 1 | 2.1 mm | 2.9 mm | 9° | 33° | 51° | 4.3 mm | Right | Right |
| Striatum 2 | 0 mm | 1 mm | 90° | 45° | 90° | 6 mm | Right | Left |
| Midbrain | 3.6 mm | 1 mm | 107° | 45° | 0° | 3.2 mm | Right | Left |

Before moving the 4-shank probes with the angle and depth parameters below, the 2D face of the 4-shank probe was positioned perpendicular to the AP (anterior-posterior) axis of the brain and above the right hemisphere of the brain. All angle and depth parameters were measured for the first shank that was closest to the sagittal suture. For striatal recordings, we used two angle strategies, (1) from the method described above and (2) parameters from Peters et al. 2021 (*37*), where the authors mapped cortical-striatal functional correspondence. AP coordinates were referenced to the bregma position, ML (medial-lateral) coordinates were referenced to the sagittal suture, azimuth angles were counterclockwise rotation around the DV(dorsal-ventral) axis from the dorsal-to-ventral view, elevation angles were clockwise rotation around the AP axis from the posterior-to-anterior view, spin angles were counterclockwise rotation around the probe axis from the probe base-to-tip view.

On or before the first day of recording, a 1-2 mm diameter craniotomy was prepared with a dental drill over the target brain area under anesthesia, leaving the dura intact. The craniotomy was protected with transparent dura-gel (Dow Corning 3-4680 Silicone Gel), with a further 3-D printed protective cap sitting on top of the recording chamber. After several hours of recovery, mice were head-fixed in the recording setup. A light shield cone was installed to connect between the recording chamber and the top edges of the ipad screens to block imaging light from the mouse’s eyes while achieving maximum probe angle flexibility. We used internal reference configuration for all recordings, with ground and reference connected. In detail, a probe was directly inserted through the dura gel without solution bath and the reference site on the probe tip was used for grounding and referencing. Prior to each insertion, the electrode shanks were coated with CM-Dil (Invitrogen), a red lipophilic dye, for later histological reconstruction of probe tracks.

### Histology

To recover electrode tracks, the brain tissue was first extracted after perfusion with 4% formaldehyde under terminal anesthesia, followed by tissue clearance with the iDISCO protocol (*81*). Image stacks of the brain sample were acquired with a light-sheet microscope (LaVision Biotec UltraScope II) at 561 nm excitation channel for DiI and 488 nm for autofluorescence. The 3D sample volume was then registered to Allen CCF (*55*) using open source package ‘ara_tools’ based on image registration suite Elastix (https://github.com/SainsburyWellcomeCentre/ara_tools). The electrode tracks were manually traced within the 3D sample volume space using open source package ‘lasagna’ (https://github.com/SainsburyWellcomeCentre/lasagna).

### Power spectrum analysis (fig. S1)

To compute the power and power ratio map in the 2-8 Hz frequency band across the cortex efficiently, we computed the power spectrum on the temporal components of the data (S*V) after singular value decomposition, and transformed the power spectrum back to the 2-dimensional space by multiplying the spatial components (U) (*49*). Power spectrum was performed using multitaper fourier transform (*82*) and averaged across multiple 20-second epochs of data. Power spectrums across mice were averaged to construct a 2-8 Hz “power map” across the cortical pixels (fig. S1C). We also used a fast Fourier transform approach to construct the 2-8 Hz “power map” and achieved the same results (data not shown). Power spectrums of example pixels were constructed with the original data, instead of SVD components (fig. S1D).

### Spiral detection and quantification (Fig. 1, fig. S1 to S9)

#### Spiral detection algorithm (Fig. 1, fig. S1, fig. S2)

To capture the dynamics of the widefield time series data, we first took the first-order derivative of the calcium traces, approximating a deconvolution to better match underlying firing rates (*75*), and then filtered the traces with a second-order Butterworth bandpass filter at 2-8 Hz with zero phase distortion. To extract the spatial distribution of the oscillation phase angle over time, the Hilbert transform is applied after filtering to each pixel independently. For visualization, the phase information for all pixels was plotted together to construct phase maps over time (Fig. 1A and B, and fig. S1).

To automatically detect spirals (fig. S2), we took advantage of the fact that phase angles along the spiral circle for any given radius cumulatively increase by 2π. We designed a three-step algorithm with a coarse-to-fine approach to identify spiral centers and radii. Step 1: Coarse grid search for spirals. Each frame is zero padded with 120 extra pixels (2 mm) on each of the four edges first to effectively detect spirals near the edge or with large radii. Spirals were first identified on a coarse grid (10 x 10 pixels, 173 x173 μm spacing). Since the phase angle distribution within a spiral is noisy at times, we defined two broad criteria for spiral detection: (1) angular differences of 10 evenly sampled points along the spiral circle cumulatively added up to near ±2π (2π ± 0.32π), with opposite signs representing clockwise and counterclockwise spiral directions; (2) at least one out of the 10 points sampled lies in each of the 4 quadrants within 0 to 2π range. At each search grid, 3 small circles (radii at 10,15, 20 pixels, which are 173, 260, 346 μm) were sampled independently. If at least 2 out of 3 sampled circles passed the two criteria above, then the grid coordinate was considered a candidate spiral position. Step 2: Clustering nearby candidate spiral centers. A robust spiral was often detected at multiple nearby candidate grid points. Spiral centers separated by a Euclidean distance less than 15 pixels (260 μm) were grouped together, and the mean coordinate of the group was taken as the intermediate candidate spiral center. Step 3: Refined spiral search and radius determination. The final positions of spirals were refined with the same algorithm from step 1 and 2 within a 20 x 20 pixels ROI at full resolution based on the intermediate candidate spiral coordinates from step 2 through a pixel-by-pixel search. The final radius was tested from 10 to 100 pixel values at 10 pixel steps (from 17.3 μm to 173 μm, at 17.3 μm steps), and the maximum radius was identified as the largest radius for which the two criteria in Step 1 were satisfied.

#### Phase randomization control for spiral occurrence (fig. S3)

Spirals can randomly occur within spatially and temporally smooth dynamics. To ensure that the spirals we observed were not by chance, we constructed surrogate data using a Fourier-based phase randomization, which preserved both the spatial and temporal correlations of the raw data (*14*, *83*). We performed the fast Fourier transform on the 3D raw data (x, y, t), randomized the phase values of the Fourier components by shuffling the index of the 3D phase matrix, and then reconstructed surrogate data with the inverse Fourier transform. To avoid including areas outside of the brain in the anterior space for the Fourier transform, we drew a rectangle in the posterior cortex and used only this area for both raw data and surrogate data comparison. We then applied the spiral detection algorithm to the raw data and surrogate data within the rectangle region, and compared distributions of detected spirals (fig. S3). Peak spiral density for spirals with radius greater than or equal to 0.69 mm within the somatosensory cortex (SSp) were significantly higher in the real data than in the surrogate data across 15 mice tested (fig. S3). Therefore, we only included spirals that had radius greater than or equal to 0.69 mm for further analysis.

#### Spiral sequences and permutation control (Fig. 1)

To group traveling spirals into spatiotemporally connected spiral sequences, we classified spirals that were less than 30 pixels (0.5 mm) apart in Euclidean distance in neighboring frames to be in the same sequence group (Fig. 1D). To determine whether the spiral sequences we observed could arise from chance with the same overall rate of spiral occurrence, we permuted the frame identity for all included spirals (those with radius greater than or equal to 0.69 mm) within a session 1000 times and recomputed the sequence grouping. Proportions of spirals that belonged to a sequence with duration lasting for more than one frame (29 ms) were consistently and significantly lower in the permutations compared to our observed data (p < 1×10^-7^,unbalanced two-way anova; Fig. 1D), while the proportion of single-frame spirals in the permuted distribution was significantly higher than in observed data (p < 1×10^-7^, student’s t-test; Fig. 1D). We therefore excluded single-frame spirals from further analysis. Nevertheless, single-frame spirals (29 ms) were also spatially concentrated within the SSp center (fig. S6).

#### Spirals in the cortical LFP activity (fig. S4)

To capture spiral waves in the cortical LFP activity without GCaMP sensors, we inserted a 4-shank Neuropixels probe with 0.7-mm span of active sites on each shank in wild type C57BL/6 mice at a shallow angle in the middle of SSp (fig. S4). The shallow angle helped us record as many SSp regions as possible in a single recording, since a steep probe angle usually captures neural activity in a single region across layers. LFP was downsampled to 35 Hz after applying a low pass filter at 17 Hz. To investigate traveling waves within the same frequency range as the widefield data in GCaMP expressing mice, we then applied a second-order Butterworth bandpass filter at 2-8 Hz and the Hilbert transform for each individual site. All sites were then rearranged as a 8×48 site matrix based on probe site location, and LFP activity map and phase maps over time were plotted (fig. S4).

#### Spiral center density estimation (Fig. 1, fig. S5, fig. S6)

To estimate the spatial density of spiral centers in cortex, we drew a local square (0.4*0.4 mm^2^) around each unique spiral and counted total spiral centers within the square. All spirals (with radius greater than or equal to 0.69 mm) that were within a spiral sequence (>= 2 frames) were included for the spiral center density plot. Given total frame numbers within a session and the sampling frequency of 35 Hz, we then converted these counts to density rates with units of spirals/(mm^2^*s). For the summary spiral density plot (Fig. 1E), all 15 sessions were concatenated for spiral density estimation.

#### Spiral symmetry quantification (Fig. 3, fig. S7)

Three peaks in the spiral center density plot were observed on each hemisphere, centered on SSp, MOp and MOs respectively (Fig. 1, fig. S7). To determine whether spirals are symmetrical and therefore are traveling in opposite directions in neighboring structures, we grouped all detected spirals into 6 regions for quantification: SSp, MOp and MOs regions in the left and right hemisphere. The borders between regions were based on Allen CCF (*55*) after alignment in Methods “Widefield imaging and data processing”. We compared spiral directions between MOp and SSp within each hemisphere (left MOp-SSp, right MOp-SSp), MOp and MOs within each hemisphere (left MOp-MOs, right MOp-MOs), and between each of the three regions across hemispheres (SSp left-right, MOp left-right, MOs left-right). When a pair of spirals were detected simultaneously in a single frame in both neighboring regions, the directions of the spirals were compared. When the two neighboring spirals were traveling in opposite directions, they were considered to be matched, in the sense that our hypothesis based on the mirrored topographic organization of these regions (*30*) predicted mirrored dynamics.

#### Synchrony index, spirality index and plane wave index (fig. S8)

To quantify traveling waves across all frames, we designed a synchrony index and a spirality index (fig. S8). Synchrony index (Rsync) is the mean resultant vector length of all phase angles in a frame (length of the vector sum/total length):

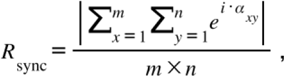

where α_x,y_ is the phase angle of the [x, y] -th pixel within a frame of frame size [m, n]. Each phase angle is represented as a vector with unit length of 1, resulting in a total length equal to the number of pixels in the frame (m x n). Synchrony index (Rsync) is bounded between 0 and 1. Synchrony index close to 0 indicates all phase angles are random and Synchrony index close to 1 indicates all phase angles are uniform.

However, the synchrony index does not differentiate spiral patterns from randomness. The synchrony index for spiral-like patterns is also close to 0, since all phases with opposing directions are represented in a spiral. Therefore we designed a spirality index (Rspiral), to quantify how close the phase map (A) from a single half-hemisphere of size [m, n/2] from a sampled frame with size [m, n], matched to a perfect spiral template (B) of the same size ([m, n/2], fig. S8). Angles (β) within the spiral template (B) were constructed with a four-quadrant inverse tangent function of the position matrix [*u*_*x*_, *v*_*y*_], which were bound in the closed interval [-π, π]:

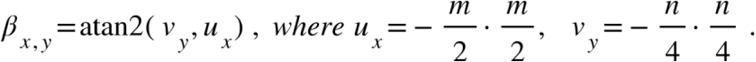

Spirality index (Rspiral) is the mean resultant vector of the angular difference map (θ):

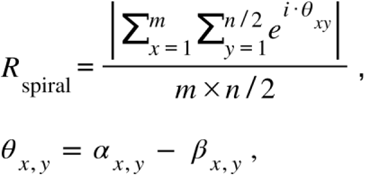

where θ is the angular difference between phase angles (α) in a sampled half-hemisphere frame (A) with [m, n/2] size and a spiral phase template (β) with a perfect spiral (B) of the same size. 0 indicates no spiral-like features in the frame, while 1 indicates a spiral-like frame.

Sum index (S) is defined as:

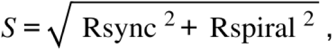

which is also bounded between 0 and 1, as frames with high synchrony index have low spirality index and vice versa (fig. S8F).

We thereafter quantified the relationship between (1) synchrony index (or spirality index) and (2) mean 2-8 Hz amplitude (or facial motion energy), by averaging the synchrony index (or spirality index) for frames within each mean 2-8 Hz amplitude (or facial motion energy) bin. The facial motion energy for each frame was calculated as the cross-pixel sum of the absolute difference between consecutive frames of the mouse face video, with higher value indicating higher motion state. Mean 2-8 Hz amplitude for each frame was calculated as the mean 2-8 Hz amplitude across all pixels within the widefield cortex boundary. The amplitude of 2-8 Hz oscillation for each pixel was the complex magnitude of the analytic signal from the Hilbert transform after second-order Butterworth bandpass filter at 2-8 Hz for each pixel trace.

To systematically quantify plane waves, we first calculated the cortical flow vector field using phase maps from consecutive frames within the 2-8 Hz range, as described in the Methods section “Traveling wave quantification with optical flow”. We then employed an order parameter, the plane wave index, to measure the coherence of the plane wave in each frame (*7*, *84*). This index is defined as the average normalized vector field velocity (φ):

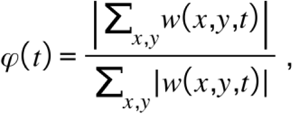

where w(x,y,t) is the flow vector field at position (x,y) in the frame at time t. Therefore, φ(t) is the absolute length of summed vector fields normalized to total absolute vector length at time t. The plane wave index is bounded between 0 and 1, with values close to 1 indicating coherent motion of all flow vectors and values close to 0 indicating random motion. We used a threshold of 0.6 to identify frames with significant plane wave activity, similar to thresholds applied in previous studies (*7*, *84*).

#### Spiral speed (fig. S9)

We calculated optical flow from sequential pairs of phase-map frames with the Horn-Schunck method (*7*, *85*) to illustrate directions of wave travel (Fig. 1C). To quantify spiral wave speed accurately (fig. S9), we sampled 12 evenly spaced points on circles of varying distance from the spiral center within each detected spiral. We used all spirals within a 1.7×3.4mm square centered on SSp in the right hemisphere across all mice (Fig. 1F, g, fig. S9). Angular speed (ω, rad/s) for any point within the spiral was calculated with the following equation:

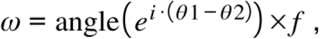

where θ1 is the phase angle for the sampled point of current frame, θ2 is the phase angle for the sampled point of the next frame, *f* is the sampling frequency (35 Hz). Angular difference is wrapped in the interval [-π, π] with Euler’s formula and angle function in matlab.

We then calculate linear speed (v):

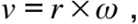

where r is the distance of the sampled point to the spiral center, ω is the angular speed for the sample point.

### Single neuron axonal arbor analysis (Fig. 2)

To investigate the morphological basis of spiral waves in the cortex, we used the Single Neuron Reconstruction Dataset (*52*), in which 1,741 neurons from various brain regions were sparsely labeled and reconstructed after whole-brain imaging. Out of 1,741 neurons throughout the entire brain, 435 neurons were labeled within the left sensory cortex. We analyzed 120 neurons within the motor cortex on the same left hemisphere separately with the same methodology (fig. S10). The sensory cortex includes visual cortex (VIS), auditory cortex (AUD), retrosplenial cortex (RSP), primary somatosensory cortex (SSp) and secondary somatosensory cortex (SSs). To reveal the local axon architecture within the sensory cortex, we excluded long-range axon projections, i.e. projections to the subcortical areas, to the contralateral hemispheres, and to motor areas of cortex. For SSs and AUD, where ipsilateral long-range projections were prevalent, only axons restricted to the local regions (SSs and AUD only) were analyzed. The singular value decomposition was computed for each neuron’s 2D point cloud (in the AP-ML plane) of axon terminals (ending points of axon branches). The first singular vector was taken as the dominant direction of the axonal arbor. A “polarity index” was computed to capture the elongation of the axonal arbor:

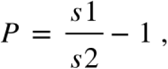

where P is the polarity index, s_1_ is the first singular value, and s_2_ is the second singular value. A polarity index closer to 0 indicates symmetrical distribution of axon terminals on the first two major axes, and larger values indicate a dominant distribution of axon terminals on the first major axis.

Axon-to-SSp-center angle (θ) is calculated as the angle between (1) the axonal first singular vector (s1) and (2) the vector from the SSp center to cell soma. The permuted distribution is generated by scrambling the soma position of each neuron 100 times while preserving the direction of the singular vector, and re-calculating the SSp-center bias.

To compare axon orientations to spirals in the mesoscale activity, we computed a mean spiral flow field by averaging the optical flow across all spirals with a large radius around the SSp center that were counterclockwise traveling in one hemisphere in all 15 mice used. We then sampled flow vectors within the flow field at the soma positions of all 435 sensory neurons used in the axon vector map (u), forming a paired flow vector map (v). We then designed an axon-flow field vector matching index, which is the sum of absolute values of the dot product of all axon vectors (u) and flow field vectors (v) at the same soma location, normalized by the sum of dot products of vector length:

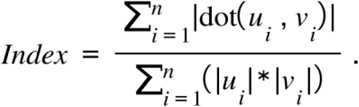

If angles between two vectors at the same soma location are ±90 degrees, then the dot product is zero; if they are matching either at 0 or 180 degrees, the absolute value of the dot product is maximized. As a result, an index of 1 indicates perfect matching of all axon and flow vectors, and an index of 0 indicates total randomness. Axon vector length is the polarity index (P) of the neuron. The null hypothesis that axon polarity is unrelated to activity flow vectors was tested by permuting the flow vectors, i.e. assigning each of the 435 spatial coordinates to have a flow vector randomly drawn without replacement from the set of all 435 flow vectors in the original map. The null distribution was generated by permuting the flow vector map 1000 times and recalculating the matching index.

### Coupled oscillators with circular connectivity bias and mirror symmetry (fig. S10)

#### Coupled oscillator model with Kuramoto equation

In order to capture spiral waves mathematically, we built a minimal model of spatially-coupled phase oscillators which could be naturally extended to include circular connectivity bias (*86*, *87*). The rate of phase change for the i-th oscillator 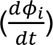 in a system composed of N oscillators is following the Kuramoto equation (*86*, *88*, *89*):

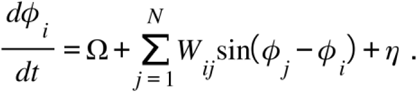

Each oscillator has an intrinsic natural frequency at Σ, and is coupled to all other oscillators with coupling weights *W*_*ij*_. The interactions between two oscillators depend sinusoidally on the phase difference between them. η is the noise term.

#### Isotropic model

We begin by generating a uniformly random point set in 2D space (fig. S10E and F), where effective coupling weight (*W*_*ij*_) for each pair of node *i* and *j* exponentially decays with their Euclidean distance *d*_*ij*_, mimicking cortical synaptic connectivity (*90*, *91*):

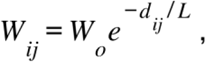

where *d*_*ij*_ is the Euclidean distance between node *i* and node *j* in the random point set, L is the effective length scale of the coupling, and *W*_*o*_ is the scale factor for the coupling matrix.

Consistent with previous results (*91*, *92*), traveling waves naturally emerged when L was small (e.g. the coupling is local) and *W*_*o*_ was sufficiently large to pull oscillators into agreement (fig. S10E). We call this model with local decaying weights (*W*_*ij*_) an isotropic model.

#### Circular bias model

To construct a model with circular connectivity bias building on top of the isotropic model above (fig. S10F and G), we defined an additional nondimensional parameter (σ) for each connected node pairs:

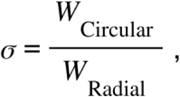

where *W*_*Circular*_ is the circular connectivity strength and *W*_*Radial*_ is the radial connectivity strength, referencing the center of the domain (2D random point space; fig. S10F). *W*_*Radial*_ points to the direction of the radial vector connecting the midpoint of the neuron pair and the center point of the 2D space, while *W*_*Circular*_ is orthogonal to *W*_*Radial*_ and thus tangential to circular node connections The coupling weight (*W*′_*ij*_) can therefore be turnably adjusted to have a strong circular bias with a higher σ value through:

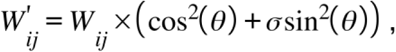

where θ is the angle between (1) the vector formed by the node pair and (2) the radial direction for the center of the node pair to the center of the domain, *W*_*ij*_ is the coupling weight defined by the Euclidean distance between node *i* and *j* in the isotropic model. The adjusted coupling weights *W*′_*ij*_ are thereafter normalized by the mean of all weights.

When sigma (σ) is greater than 1, there is an azimuthal/circular bias; if equal to 1, the connectivity is isotropic; and when less than 1, the connectivity has a radial bias.

#### Spirality index

We sought to characterize the presence of spiral waves in the baseline model and circular bias model for topographically coupled oscillators (fig. S10F and G). As a complementary tool to study the presence of spiral waves, we leveraged a population scale metric related to synchronization to look for the presence or absence of spirality (*Sp*), which measures how closely a phase map matches a perfectly centered spiral. This is the same as the definition of Spirality index (Rspiral) as described in Methods section “Synchrony and spirality index” (fig. S8):

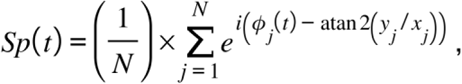

where *Sp*(*t*) is the spirality at time *t*, [*x*_*j*_, *y*_*j*_] is the position index for the *j*-th node in the 2D random point set, φ_*j*_(*t*) is the phase value of the *j*-th node at time *t* of simulation, *ata*φ2(*y*_*i*_/*x*_*i*_) is the phase value of the j-th node of the center spiral template, and N is the total number of nodes in the 2D random point set.

Spirality (*Sp*) measures how close the current configuration is to the perfect spiral configuration centered at the center of the field of view on a scale of 0 to 1, with 1 only being achieved when the full field is a spiral centered in the field of view. Equipped with this tool, we sought to characterize spiral waves in our baseline model. Across 1000 simulations (robust against 10 random seeds), we found that spiral waves were more likely near the transition between a desynchronized regime and a synchronized regime. In the crossover vicinity (reflected by noise and scale of coupling critical line separating these two regimes), we observed a dramatic uptick in the spirality of these fields (fig. S10F and G). We interpret this result as consistent with the proliferation of cortices/defects in a continuous spin-system as it undergoes a BKT phase transition (*93*, *94*). In the desynchronized regime, there were many defects (including spiral waves) with very short lifetimes as they bounced around and annihilated each other under noisy driving. In the synchronized regime, the dominant defect was the spiral wave because they were topologically protected against continuous deformations leaving them only to find an opposite spiral to annihilate or leave/enter through the boundary of the domain.

#### Comparison of isotropic model and circular bias model

To compare the isotropic model and circular bias model, we first defined the circular connectivity bias σ parameter at 50 for circular bias model (σ at 1 for isotropic model by default) and explored the spirality index change as a function of weight scale *W*_*o*_ and noise η (fig. S10G). We used a total of 1024 oscillators (32^2^). The coupling length scale L was set to 0.05, corresponding to a 63% decrease in coupling strength at a 50 µm distance from a given cell in a network scaled to 1 mm in width or height (or 5%). Coupling weights of the network increased linearly by dialing up the coupling weight scale *W*_*o*_ between 0 and 1 along the X-axis of the phase space sweep (fig. S10G). The noise term η was sampled from gaussian distribution, applied independently to each oscillator. The Y-axis of the phase space sweep (fig. S10G) corresponds to the scale of the standard deviation of this noise.

In addition, we investigated the spirality index as a function of weight scale *W*_*o*_ and sigma σvalues, without noise. We chose initial seeds where spirals were present but not centered in the isotropic model (sigma σ = 1). Spirality increased as a function of sigma in the circular bias model (sigma > 1), indicating circular connectivity helped spirals stabilize in the center of the circular connectivity network (fig. S10H). In seeds where spirals were already in the center of the 2d space, increasing sigma had minimal effect on spirality, as the spirality index is maximized to begin with.

#### Mirror symmetry model

To explore how mirror symmetry (Fig. 3) affects the spiral distribution, we constructed a mirror symmetry model (fig. S10I). We filled 4000 cells in the bottom right quadrant of the 2d space first with random normal distribution, with effective coupling weight between each cell pair defined as an exponential decay function proportional to the Euclidean distance between them (same as in isotropic model). We then flipped a random quarter of cells to one of the remaining three quadrants of the 2d space (repeated 3 x), and kept the cell-pair weights defined before flip. By doing so, we created a mirror symmetry model, where the cell positions and connectivity weights were mirrored in neighboring quadrants (cell pairs that were close in space before flip were still tightly coupled after flip, despite flipped quadrant position), while local connectivity within a single quadrant was still preserved. To construct a non-symmetrical model in comparison to the mirror symmetry model (fig. S10I), we kept the cell positions after the flip in the mirror symmetry model, but re-calculated the cell-pair weights with exponential decaying function across all cells (no quadrant symmetry). Weight matrix was normalized by the mean weights of all cell pairs in both isotropic model and mirror symmetry model, for a fair comparison. We simulated 150 random seeds with the same initial condition for both models. Spiral centers were detected in each simulation seed with a spiral detection algorithm as in widefield imaging data, and concatenated across 150 seeds for comparison. Spiral density was normalized to the same peak density value for model comparison. An absence of spiral centers along the borders in the mirror symmetry model was observed, as in the widefield imaging data.

### Linear regression within cortex (Fig. 3, fig. S11)

Reduced rank regression was applied to predict the posterior sensory cortical activity from the anterior motor cortex, and to predict the right hemisphere from the left hemisphere. Reduced rank regression has the advantage of finding the subspace of regressor activity that is maximally predictive of the target activity, thus acting as both a regularization and a characterization of dimensionality (*95*). To divide the anterior/posterior cortex as regressor and target, widefield data (X*Y*t) was first registered to allen brain atlas CCF coordinates (*55*), and divided into anterior cortical data and posterior cortical data along the border of SSp and MOp respecting the connectivity reciprocity (*30*) and spiral symmetry we observed. To divide the left/right cortex, the midline was chosen as the dividing line after registration. The first 20-minutes of the widefield data in each session was used as training data, and a 10-minute data section following the training data was used as testing data for cross validation. To account for data collinearity, we applied singular value decomposition (D = USV^T^) on the regressor data and used the top 50 temporal components (S*V) as the regressor in the regression model, which accounted for 97.3 ± 0.5% (mean ± SEM, 15 mice) of total variance before compression, as previously stated in Method “Widefield imaging and data processing” section. The target data was kept in the original data space (pixels*t). Prediction accuracy was calculated using Coefficient of determination (R^2^) on the testing dataset:

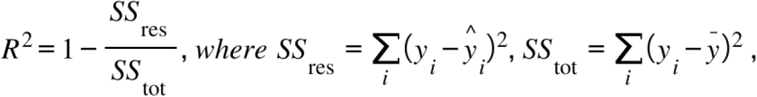

where *SS*_*res*_ is the residual sum of squares, *SS*_*tot*_ is the total sum of squares, *y*_*i*_ is the i-th value in the observed data, *y*^_*i*_ is the predicted data, *ȳ* is the mean of the observed data.

To visualize the regression kernel representation in the cortex space, the regression kernel was first transformed back to the pixel space. Kernel maps for example pixels were presented as different color plots overlaid with transparency scaled by kernel weights (Fig. 3D to G, H, J, and fig. S11). Example regressor pixels in Fig. 3D to G were chosen as the pixel with maximum value in the weight matrix.

### Bulk axon projection map and matching index (Fig. 3)

To visualize the long range axon projection maps from sensory cortex to the contralateral hemisphere, or to the motor cortex, we selected eight viral tracer injection experiments targeting eight different sensory modalities within the sensory cortex from the Allen Mouse Brain Connectivity Atlas (*35*). The same session IDs were used to identify Neuropixels targeting angles and positions in the subcortical areas in Methods “Simultaneous Neuropixels recording and widefield imaging” section. To combine axon projection maps together, intensity values for injection sites in the sensory cortex were masked first and then normalized across sessions to account for intensity differences between injection site and target axon region, as well as differences across sessions respectively.

To quantify whether the activity kernel patterns and axon projection patterns matched, we designed a pattern matching index (PIndex) which is the sum of matrix dot product between activity kernel and axon projection map of the same size across all 8 different sampled sensory areas:

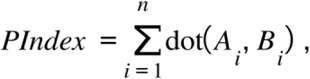

where *A*_*i*_ is the activity kernel for one of the n (n = 8) sampled sensory areas, *B*_*i*_ is the axon projection map for one of the 8 sampled sensory areas.

We used a single common axon projection session for each of 8 sensory areas to compute the matching index, together with mean activity maps across 15 mice. Small variations were observed between the activity map and the axon projection map due to variations of brain registration for each mouse, as well as the slight difference of sampling points for activity map and projection maps. However, the general patterns of both maps matched well for left-to-right hemispheres (Fig. 3H and I) and motor-to-sensory cortex (Fig. 3J and K), compared to permutation. Permutation was performed by randomizing the identity of the 8 axon projection maps 1000 times. For visualization purposes, we normalized pattern matching index in real data and permutation by the minimum and maximum values in the permutation set and real data together, ensuring all values are within [0,1] index scale.

### Spiral prediction from subcortical spiking activity (Fig. 4, fig. S12)

Kilosort2 (*96*) was used for spike sorting of the electrophysiology data from 4-shank Neuropixels recordings. Single neurons were manually curated through Phy (https://github.com/cortex-lab/phy), with violation of refractory period in spike time auto-correlogram as the main criteria for selection. Spikes of each single neuron were then binned at the widefield imaging sampling frequency (35Hz) to match the time series of the imaging data.

To predict cortical widefield data from subcortical spike data (Fig. 4B and C), time series of spiking neurons were used to predict the derivative of top 50 temporal components of the cortical widefield data. To predict the full length of the widefield data from subcortical spiking activity, we used 2-fold cross-validation. In detail, we first split the data into 10 equal-length epochs. Then we used the concatenated 5 odd-numbered epochs to predict the held-out even epochs, and vice versa. The mean variance explained map was calculated as the average of the two predictions. To account for the latency between cortical widefield data and spike data, time-shifted spiking series from -17 to 17 samples (-0.5 to 0.5 s at ∼0.03 s time steps) were also concatenated into the regressor matrix. For quality control purposes, only sessions with mean variance explained exceeding 10% were included for further analysis (29/46 sessions).

To compare spirals in the raw and predicted cortical activity (Fig. 4D, E and G), we took the derivative of the temporal components of the raw and predicted data, and reconstructed the data in pixel space (X*Y*t) after a bandpass 2-8 Hz filter. We then detected spirals in both raw and predicted cortical data by applying the automated spiral detection algorithm. Spirals in the raw cortical activity with significant radius (greater than or equal to 0.69 mm) and duration (longer than 29 ms), were considered for spiral direction comparison. Selection criteria were not applied to the predicted spirals, to maximize the numbers of corresponding spirals included in the subsequent matching test. For each spiral in the raw cortical activity, if only one corresponding spiral was found within 1 mm radius in the predicted cortical activity in the same frame, then the pair of spirals was considered as a candidate pair. We then compared the spiral directions (clockwise or counterclockwise) for all pairs of candidate spirals, and computed the spiral matching probability as the ratio of matching spiral pairs out of all candidate spiral pairs. Here, ‘matching’ means that the raw and predicted spirals share the same direction. To test the null hypothesis that neurons in the subcortical area did not share spatiotemporal relationships with the cortical data, and that matching spirals therefore only arose by chance, we computed the matching probability with permuted spiking data. We first predicted cortical activity by using subcortical spiking activity with permuted neuron identities during linear regression prediction without retraining the model, and then applied the same automated spiral detection algorithm for the permuted prediction. We then computed the spiral matching probability between the raw and permuted prediction as control.

### Traveling wave quantification with optical flow (Fig. 4, fig. S12, fig. S13)

To quantify traveling waves within the cortical widefield activity (Fig. 4, fig. S12), we estimated instantaneous velocity for each pixel in the cortex using phase information from nearby two frames. In detail, the derivative of the widefield data was first bandpass Butterworth filtered within 2-8 Hz, and the Hilbert transform was then applied to each pixel independently to extract phase information at each time point to construct the phase maps. We then applied the Horn-Schunck method (*85*) to extract optical flow from the phase maps of adjacent frames, which used an optimization approach to estimate instantaneous velocity with a global constraint of smoothness.

To compare phase maps and traveling waves of the raw and predicted cortical activity from subcortical spiking activity (Fig. 4H and I, fig. S12, fig. S13), we designed frame-to-frame “ phase matching index” and “traveling wave matching index”. First, the angular differences of the phase values or the optical flow vectors between raw and predicted data were calculated for pixels in each frame with frame size [m,n]. Phase or traveling wave matching index (M) is the mean resultant vector length of the angular difference map in a frame:

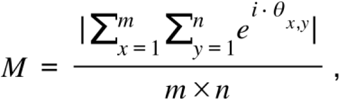

where θ_x,y_ is the phase angular differences of the [x, y]-th pixel between a raw and predicted frame. M index is bound between 0 and 1. M close to 0 indicates all angular difference values are randomly distributed and not matching, while M close to 1 indicates all angular difference values are uniformly distributed and matching.

To summarize the observation that phase or traveling wave matching index increased as mean 2-8 Hz amplitude in the cortex increased within a session (Fig. 4H and I, and fig. S13), we binned frames based on the mean 2-8 Hz amplitude and calculated the mean matching index within each amplitude bin. Only pixels with cross-validated variance explained greater than 0.1 were included in the final phase or wave matching index calculation. To construct the null distribution, we employed the same strategy as described above for spiral matching, by randomizing the subcortical neuron identities in the testing dataset for regression prediction 20 times, without re-training the model. Phase and wave matching index were then calculated between the raw phase (or traveling waves) and permuted predictions, as control distributions.

## Supplementary Figures

**Fig. S1.**
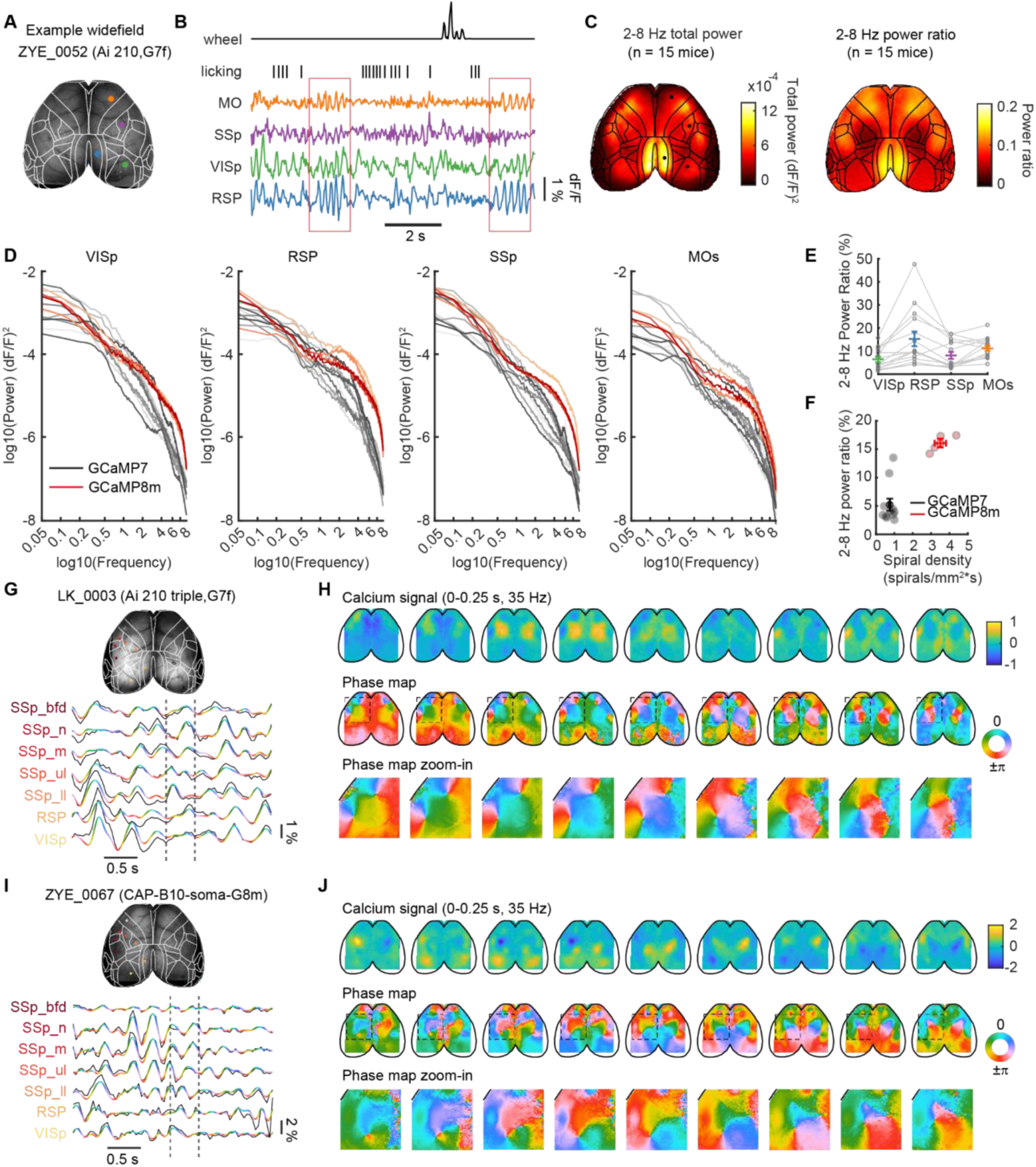
2-8 Hz power is globally distributed within the cortex and spirals pervades the mouse cortex with different transgenic strategies. (**A**) Example dorsal surface of the mouse cortex for widefield imaging in a mouse with GCaMP7f expression (ZYE_0052, Ai210 mouse injected with PHP.eB-Cre virus). The brain surface was registered to the CCF atlas (*55*), and outlines of cortical regions superimposed. (**B**) Example calcium traces from an awake, behaving mouse, with pixel locations highlighted in A. Randomly timed rewards were delivered, provoking licking and ensuring wakefulness. Two epochs of neural activity with strong 2-8 Hz oscillations between lick periods are highlighted in red boxes. (**C**) Left: cortical map of 2-8 Hz total power, averaged across 15 animals. Units: dF/F^2^. Right, cortical map of 2-8 Hz power ratio (2-8 Hz power/0.5-8 Hz power), averaged across 15 animals. (**D**) Power spectrum of example cortical regions (VISp, RSP, SSp and MOs). Each trace represents the power spectrum from a single subject. Grays: GCaMP7 mice; Reds: GCaMP8m mice. Power spectrums were computed by averaging across multiple 20s-epochs from the entire recording session of 30-60 minutes. (**E**) 2-8 Hz power ratio in 4 cortical regions. SSp had an intermediate level of 2-8 Hz signal, compared to the other three selected areas (SSp: 8.1% ± 1.5%; RSP: 15.3% ± 3.2%; MOs: 11.2% ± 1.2%; VISp: 6.4% ± 1.2%; mean ± SEM; n = 15 mice). Power ratios were calculated from example traces in D. (**F**) GCaMP8m mice had higher 2-8 Hz power ratio and spiral density compared to GCaMP7 mice, likely due to improved temporal kinetics. Gray: GCaMP7 mice; red: GCaMP8 mice. (**G**) Example calcium traces in a mouse with GCaMP7f expression (LK_0003, Ai210 triple transgenic mouse). Raw traces (black) are overlaid with 2-8 Hz filtered traces color coded by phase values after the Hilbert transform. (**H**) 9 example consecutive frames (0.25 s) of calcium signal maps (top), phase maps (middle) and spiral zoom-in (bottom) within the dashed line in G. (**I**) Same as in G, but for a mouse with soma-targeted GCaMP8m expression (ZYE_0067, wild-type mouse injected with CAP-B10-soma-GCaMP8m). (**J**) 9 example consecutive frames (0.25 s) of calcium signal maps (top), phase maps (middle) and spiral zoom-in (bottom) within the dashed line in I.

**Fig. S2.**
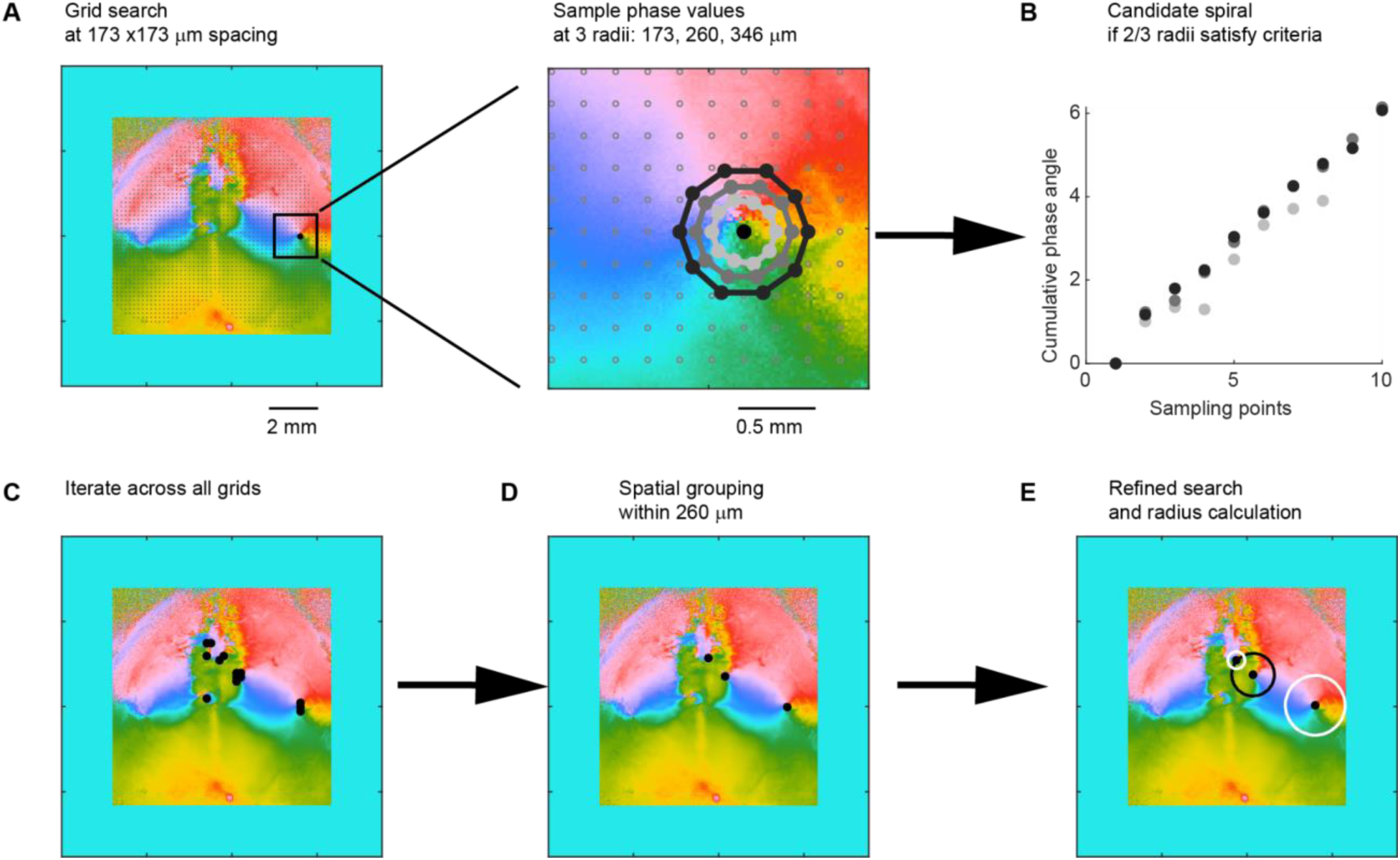
Spiral detection algorithm. A three-step process was used to identify spirals, first using a coarse grid search to identify candidate spiral centers, second by clustering nearby candidates, and third by performing a fine search to precisely identify the spiral center and radius. (**A**) Phase map frames were zero-padded at the edges and search grids (10×10 pixels, 173 x 173 μm spacing) were drawn on top of the frame within the brain ROI. At each search grid, 3 small circles were drawn (radius at 10, 15, 20 pixels, which are 173, 260, 346 μm). Within each circle, 10 phase values at equal angular spacing (from 0 to 9/5*π at π/5 steps) were sampled. (**B**) If the absolute cumulative angular difference from the first to the last sample was close to 2π (2*π ± 0.32π) and there was at least one sample at each of the 4 quadrants of the circular phase angles, then that circle was spiral-like. A grid point was a candidate spiral center if at least 2 out of 3 sampled circles were spiral-like. (**C**) Candidate spiral search was iterated across all grids. Black dots represent all candidate spiral centers that were detected in a single frame. (**D**) A robust spiral was often detected at multiple nearby grids. Spiral centers that were located within 15 pixels (260 μm) Euclidean distance were grouped together, and the mean location of the spirals within a single cluster was taken as the starting point for a final coarse-to-fine search in the next step. (**E**) The precise location of the spiral was then determined by a final refined search approach. A 21×21 pixel ROI centering the spiral candidate (grid points) was extracted, step A to C was repeated pixel-by-pixel, and final spiral center location was determined. Spiral radius was then determined by the maximum radius that satisfied criteria in B. If the cumulative angular difference from the first to the last sample was positive, then the spiral was counterclockwise traveling (white circle); otherwise, clockwise traveling (black circle). Spirals with radius less than 0.69 mm (the smallest spiral in E as an example) were excluded in the final analysis, as their distribution did not pass permutation tests and were more likely to be noise (fig. S3).

**Fig. S3.**
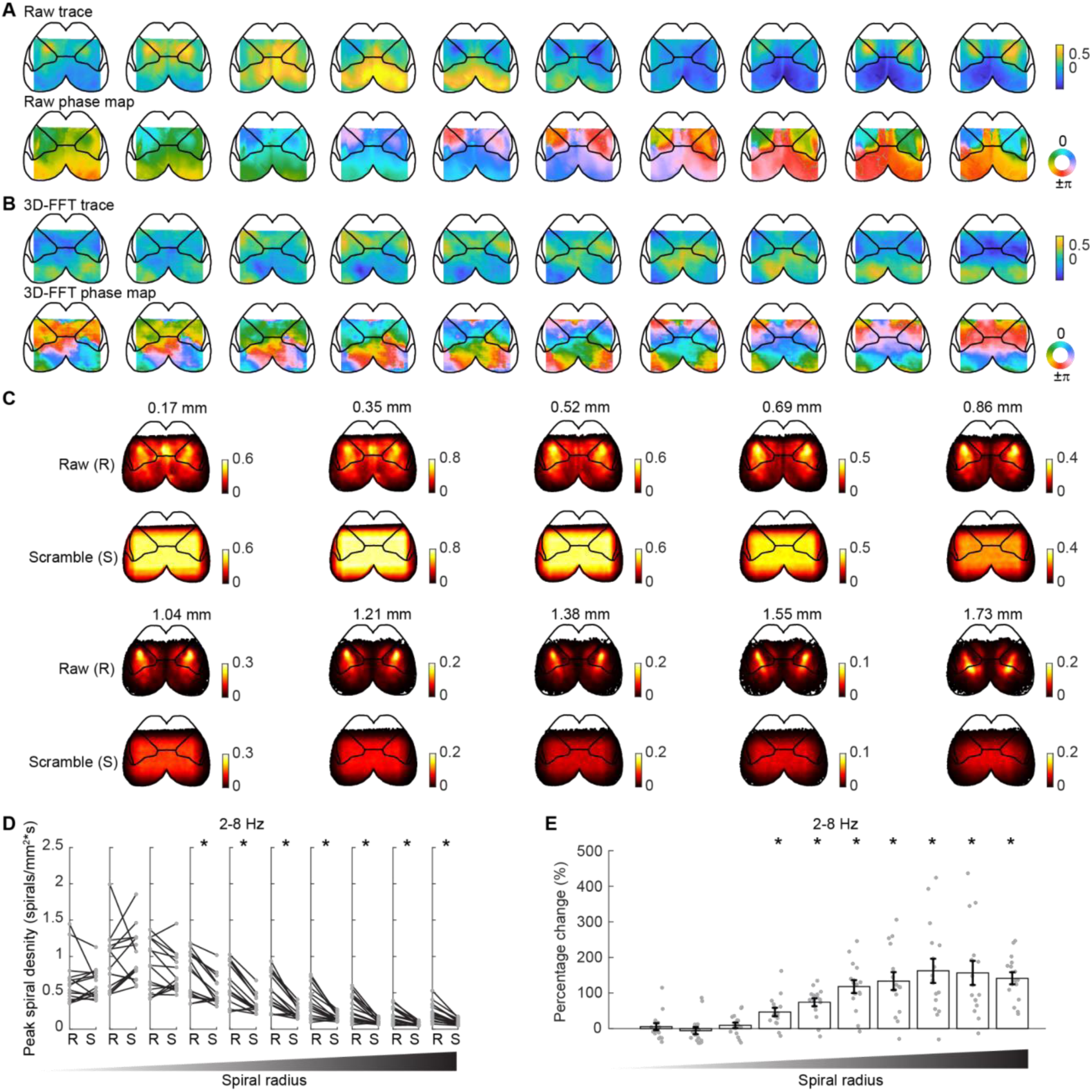
Peak spiral density exceeded fourier phase shuffled control for spirals greater than or equal to 0.69 mm radius. Spirals could occur randomly within spatiotemporally correlated data. To evaluate whether the spirals we observed within the cortex were above chance level, we constructed shuffled Fourier phase surrogate data through the 3D fast Fourier transform (FFT) from raw data (x,y,t). We shuffled the phase values of the Fourier components, and created surrogate data by the inverse Fourier transform of the components from raw data with shuffled phases. By doing this, we were able to preserve the spatial and temporal autocorrelation (or power spectrum) of the raw data. We then applied the same spiral detection algorithm to surrogate data. Spiral distributions were then compared between raw data and surrogate data across sessions. (**A**) Example sequence of widefield raw trace (top row) and phase map (bottom row). To avoid including the field of view in FFT analysis that was outside of the brain, we drew a rectangle square to cover most of the posterior brain, but avoided the anterior brain. (**B**) Sequence of surrogate data trace (top row) and phase map (bottom row). Surrogate data was constructed with 3D-FFT phase shuffling from the same section of data in A. (**C**) Spiral density plots across 10 different spiral radius bins (from 0.17 mm to 1.73 mm at 0.17 mm steps). Top rows: All spirals detected and concatenated from raw data across 15 mice. Bottom rows: All spirals detected and concatenated from surrogate data across 15 mice. Density plots from raw data and phase scrambled data for the same radius were plotted at the same color scale for comparison. Spirals were highly concentrated within the center of SSp across all radius bins. (**D**) Peak spiral density values within the SSp were compared between raw data and surrogate data across 15 mice for all 10 radius bins (0.17 mm:0.17 mm:1.73 mm). While peak density values for the first 3 small radius bins (0.17 mm, 0.35 mm, 0.52 mm) in the raw data (R) were not significantly different from the surrogate data (S), peak spiral density values significantly exceeded the surrogate data for larger radius bins (from 0.69 mm to 1.73 mm, at 0.17 mm steps). Each dot represents a single session from each of 15 mice. (**E**), Peak spiral density values were compared between raw data and surrogate data with percentage change. Spiral density significantly increased from 46.7 ± 11.7% to 141.3 ± 16.9%, for spiral radius from 0.69 mm to 1.73 mm. Paired student’s t-test, *p* = 0.001 at 0.69 mm radius, *p* = 8 x 10^-7^ at 1.73 mm radius. Note the statistical tests and results are the same for D and E.

**Fig. S4.**
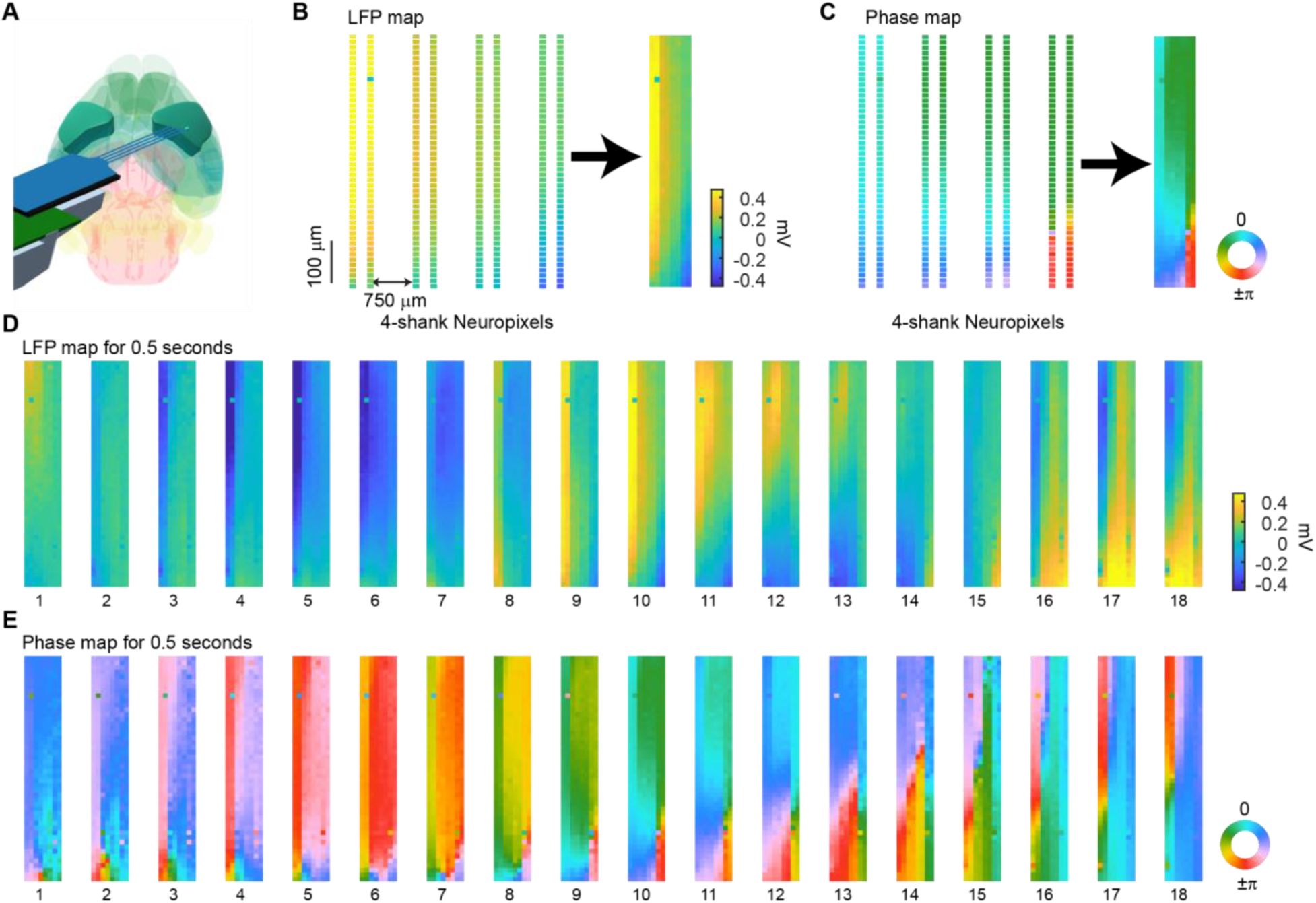
Spirals in LFP activity with 4 shank Neuropixels recordings in the sensory cortex of a wide type C57BL/6 mouse. (**A**) A 4-shank Neuropixels probe was programmed to be double-column per shank with 48 sites each column spanning 0.7 mm in length, and was inserted in a shallow angle to cover a large area of the sensory cortex in a wild type C57BL/6 mouse. (**B**) LFP amplitude of the 4-shank Neuropixels probe was filtered at 2-8 Hz and downsampled to 35 Hz. Probe columns were compressed to a 2d map for visualization. The probe dimension with 4-shanks was 0.7 mm depth x 2.25 mm width. (**C**) Phase values of filtered LFP trace at each site were extracted with the Hilbert transform, and phase map was constructed. (**D**) An example of 0.5s LFP map. (**E**) Phase map of 0.5s LFP activity in D. Note that spirals were observed in multiple frames of the 0.5s sequence.

**Fig. S5.**
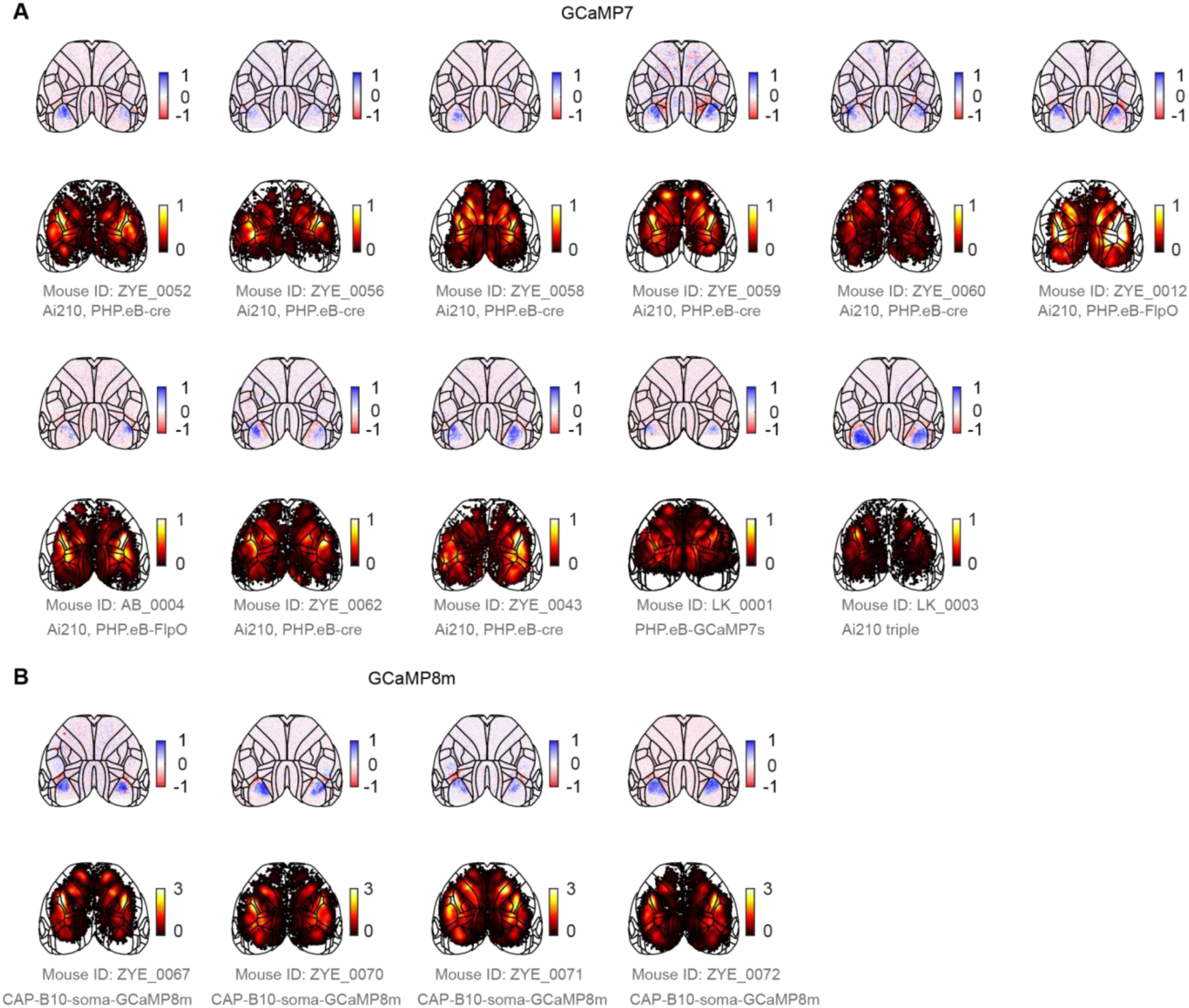
Brain registration and spiral distribution across all GCaMP7 and GCaMP8m mice. (A) Top: Brain atlas registration with visual field sign map as reference. Color bar: sign values within [-1, 1] range. Bottom: Spiral center density for all 11 GCaMP7 mice. Units: spirals/(mm^2^*s). (B) Same as A, but for 4 soma-GCaMP8m mice. GCaMP8m mice had significantly higher peak spiral rate at the center of SSp in comparison to GCaMP7 mice (GCaMP7: 0.8 ± 0.2 spirals/(mm^2^*s), mean ± STD, 11 mice; GCaMP8m: 3.5 ± 0.6 spirals/(mm^2^*s), 4 mice; p = 7.7× 10^-9^,two-sample student’s t-test).

**Fig. S6.**
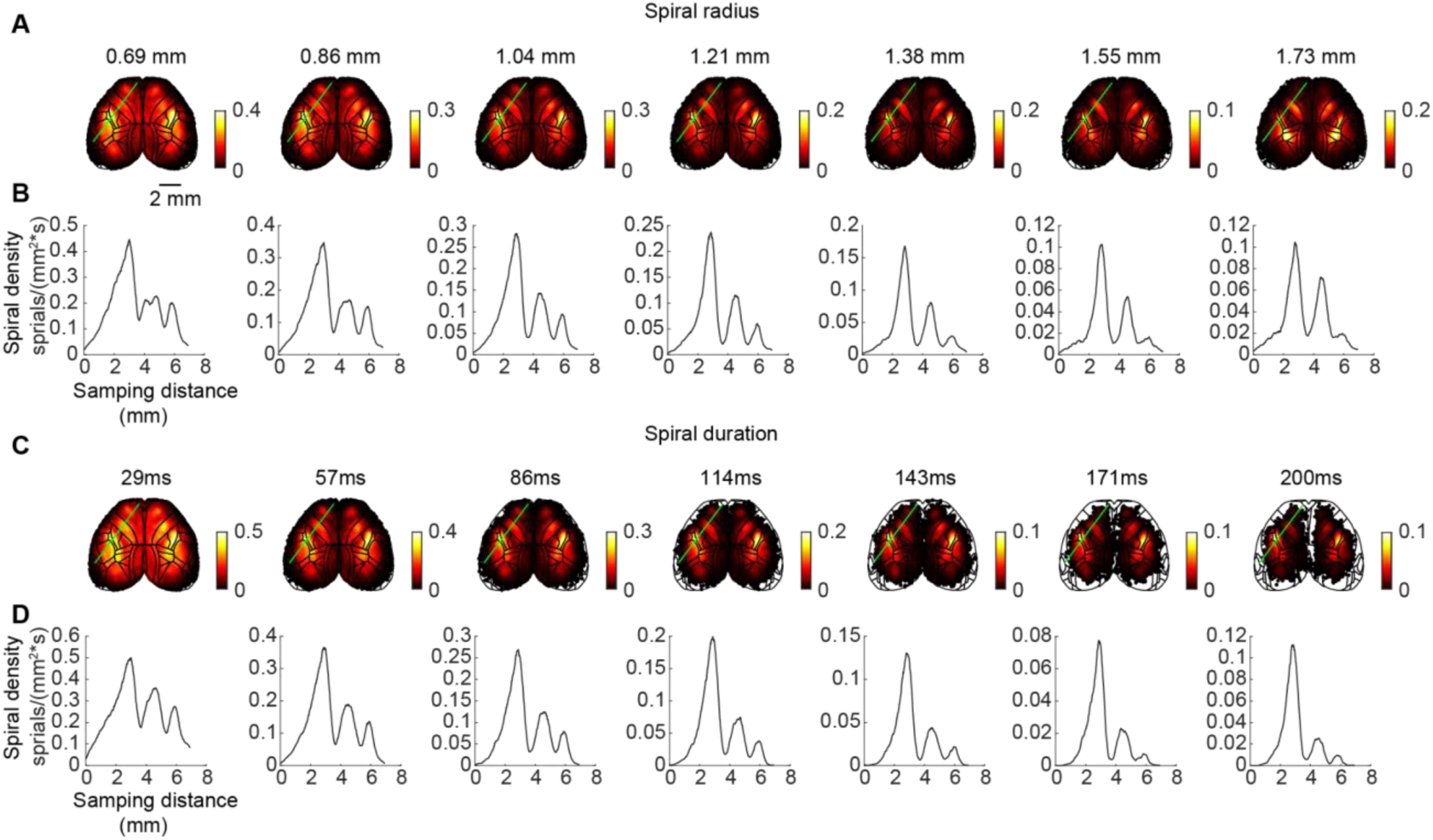
Consistent spiral distribution at different spiral radius and duration. (**A**) All spirals from 15 mice were combined and sorted into different radius bins from 0.69 mm to 1.73 mm at 0.17 mm steps. The spiral density peaks at the center of SSp were consistent across different spiral radii. The separation of spiral peaks between SSp, MOp, and MOs were prominent across spiral radius bins. (**B**) All spirals from 15 mice were combined and sorted into different duration bins from 28.6 ms to 200 ms at 28.6 ms steps. The spiral density peaks at the center of SSp, MOp, and MOs were consistent across spirals that fell within different sequence durations.

**Fig. S7.**
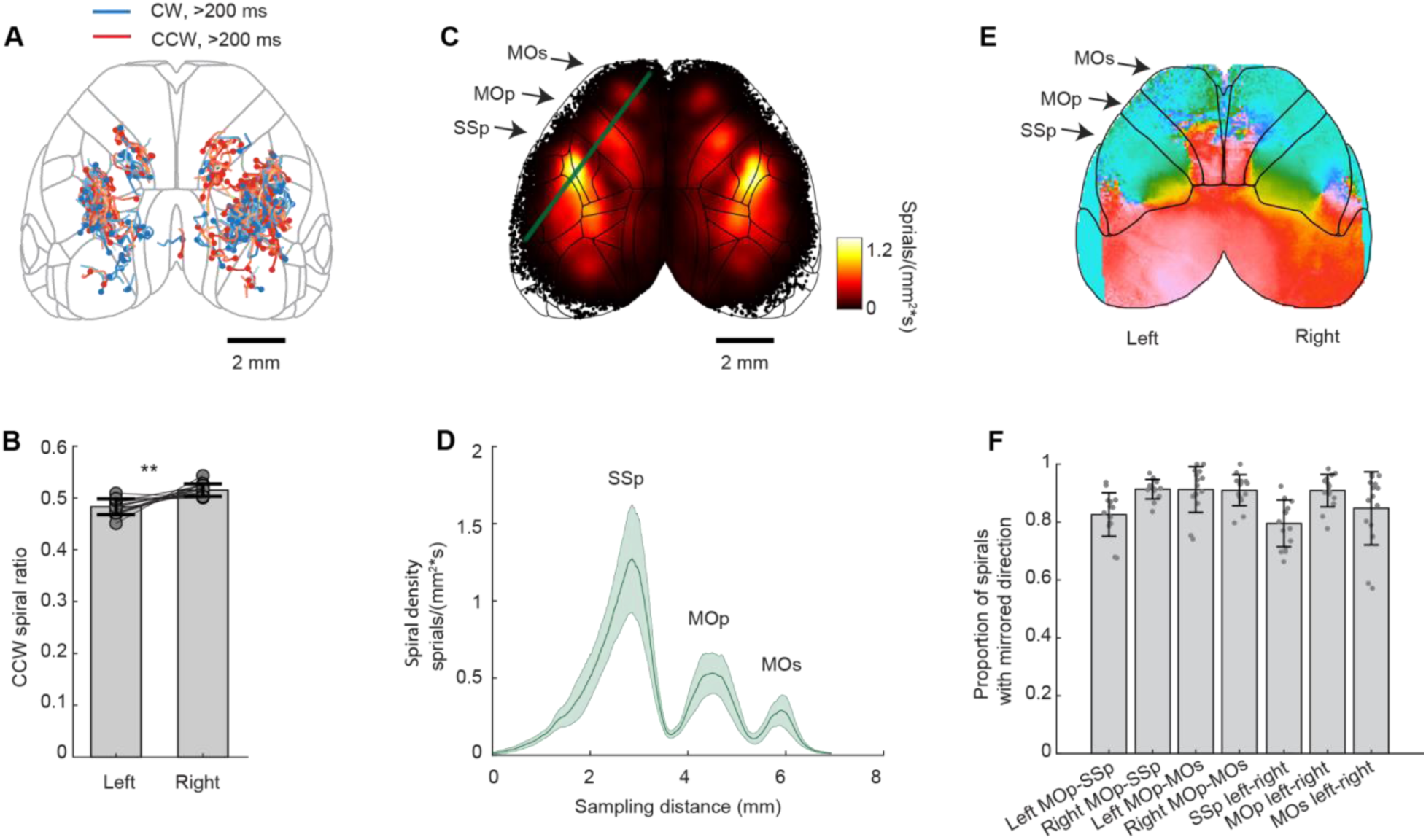
Spiral traveling direction and distribution symmetry. (**A**) Spiral centers smoothly drift over space and time. Example spiral center sequences that lasted longer than 200 ms in one example session are overlaid on the brain surface. Blue indicates clockwise traveling, red indicates counterclockwise traveling. The starting position of the sequence is indicated as a solid circle. (**B**) Spiral sequences can travel both clockwise and counterclockwise. Spirals in the right hemisphere were slightly but significantly more likely to be counterclockwise traveling than spirals in the left hemisphere across mice. CCW spiral ratio in the left hemisphere: 0.46 ± 0.04 (mean ± STD); CCW spiral ratio in the right hemisphere: 0.54 ± 0.04 (mean ± STD); n = 15 mice, Student’s t-test, p = 1.5 x10^-3^. (**C**) Spiral density distribution, summarized across all 15 mice. A line was drawn (green) to sample spiral density values across the SSp, MOp and MOs in the left hemisphere. (**D**) Spiral density at green sampling points in c illustrates 3 peaks of spirals, centered in the SSp, MOp and MOs, consistent across mice (peak density at 1.3 ± 0.3 spirals/ (mm^2^*s), n = 15 mice, mean ± SEM). (**E**) Example spiral frame with region outline overlaid. (**F**) Spirals detected on neighbor mirroring structures are mostly traveling in opposite directions. Each dot represents a session from each of 15 mice. Proportion of spirals with mirrored direction is the proportion of total frames with a pair of opposite spirals out of total frames with a candidate pair of spirals on mirroring structures. Left MOp-SSp: 82.6 ± 7.5% (mean ± STD); Right MOp-SSp: 91.3 ± 3.4%; Left MOp-Mos: 91.2 ± 7.9%; Right MOp-MOs: 91.0 ± 5.4% SSp left-right: 80.0 ± 8.0%; MOp left-right: 90.9 ± 5.6%; MOs left-right: 84.7 ± 12.7% (n = 15 mice).

**Fig. S8.**
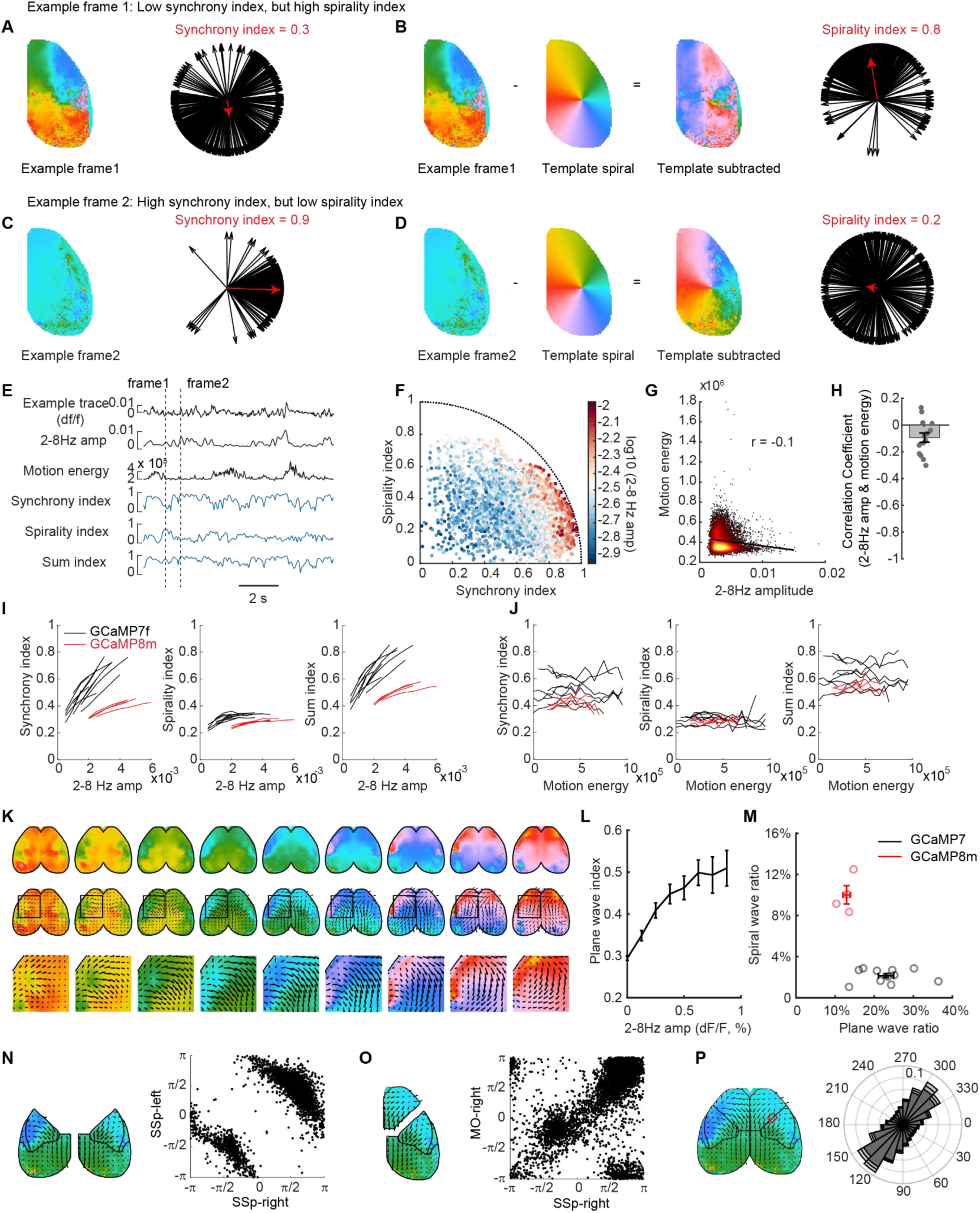
Phase synchrony, spirality index and plane wave characterization. (**A**) Phase values across pixels in the example frame 1 had close to evenly distributed phase values from -to π, which resulted in a synchrony index close to 0. (**B**) However, for the same frame 1, while its synchrony index was close to 0, it had a spirality index closer to 1, indicating it had a spiral-like pattern. (**C**),(**D**) Example frame 2 has a synchrony index close to 1, but a spirality index close to 0. (**E**) Time series of an example widefield cortical activity trace, mean 2-8 Hz amplitude, motion energy of the mouse face recording, synchrony index, spirality index and sum index for the right hemisphere of an example recording session (see Methods). Sum index = 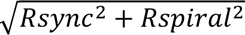, which is also bounded between 0 and 1. Note that frames that had high synchrony index often had low spirality index, vice versa. Frame examples 1 and 2 in A TO D are highlighted in the time series with dashed lines. Facial motion energy was the cross-pixel sum of absolute difference between consecutive frames of the mouse face video. (**F**) Scatter plot of synchrony index and spirality index for each frame of a 100s epoch in the same session as in E. Each dot is color coded by the mean 2-8 Hz amplitude frame-to-frame. Frames with high mean 2-8 Hz oscillation amplitude had either high synchrony index or high spirality index. However, the distance of the dot to origin (sum index) is below 1, as shown with the dashed circular boundary. (**G**) Scatter plot of 2-8 Hz amplitude and facial motion energy of an example session. 2-8 Hz amplitude was slightly negatively correlated with facial motion energy (r = -0.1). (**H**) Pearson correlation coefficient between 2-8 Hz amplitude and facial motion energy across 15 mice (r = -0.1 ± 0.04, Mean ± SEM). (**I**) Mean synchrony index (left), spirality index (middle) and sum index (right) at different 2-8 Hz oscillation amplitude bins for a random 100s epoch across 15 mice. All index values increased as 2-8 Hz oscillation amplitude increased. Distributions of 2-8 Hz amplitude in GCaMP8m mice exhibited higher 2-8 Hz amplitude overall in comparison to GCaMP7 mice, presumably due to faster rise and decay kinetics resolving 2-8 Hz range oscillation better. (**J**) Mean synchrony index (left), spirality index (middle) and sum index (right) at different motion energy bins of the mouse face recordings. Note there is no tendency of change of index across different motion energy bins, in comparison to 2-8 Hz amplitude bins in I, likely due to the weak correlations between 2-8 Hz amplitude and facial motion energy as shown in G, H. (**K**) An example plane-wave sequence across the dorsal surface of the cortex (top panel), with flow vector field overlaid (middle panel) and zoom-in of the plane wave (bottom panel). (**L**) Plane wave index (Methods) increased as the 2-8 Hz oscillation amplitude increased. (**M**) Comparison of plane wave/ spiral wave ratio across different GCaMP mice. Black, GCaMP7 mice; red, GCaMP8m mice with PHP.eB.AAV virus injection. As the kinetics of calcium sensors improved, spiral wave ratio significantly increased, while plane wave ratio decreased. (**N**) Plane waves generally respect the anatomical symmetry, like spiral waves. Illustration (left) and scatter plot (right) of mean plane-wave angles for left and right sensory cortex. Each dot represents a frame that had significant plane waves in both hemispheres. Frames across all mice were compiled, but only a random fraction (25%) are shown for clarity. The mean angles of plane waves were symmetrical across the midline, with the absolute mean angular values from the left and right hemispheres often summing to π. (**O**) Same as N, but for the motor and sensory cortex. The mean angles of plane waves were positively correlated, likely due to the dominance of waves in the anterior-posterior direction. (**P**) Plane waves respect the boundary between motor and sensory cortex. Illustration (left) and polar histogram (right) of mean plane-wave angle distribution for a square patch of pixels on the somatosensory and motor cortex border. Only frames with significant plane waves were included. Although some plane waves traveled across the boundary between the somatosensory and motor cortices, such occurrences were rare compared to plane waves traveling along the boundary (along the border at 135°, 8.8% ± 0.5%; across the border at 45°, 2.3% ± 0.2%; mean ± SEM; *p* = 3.2 x 10^-7^, paired student’s t-test). Bar edges represent mean ± SEM across 15 mice.

**Fig. S9.**
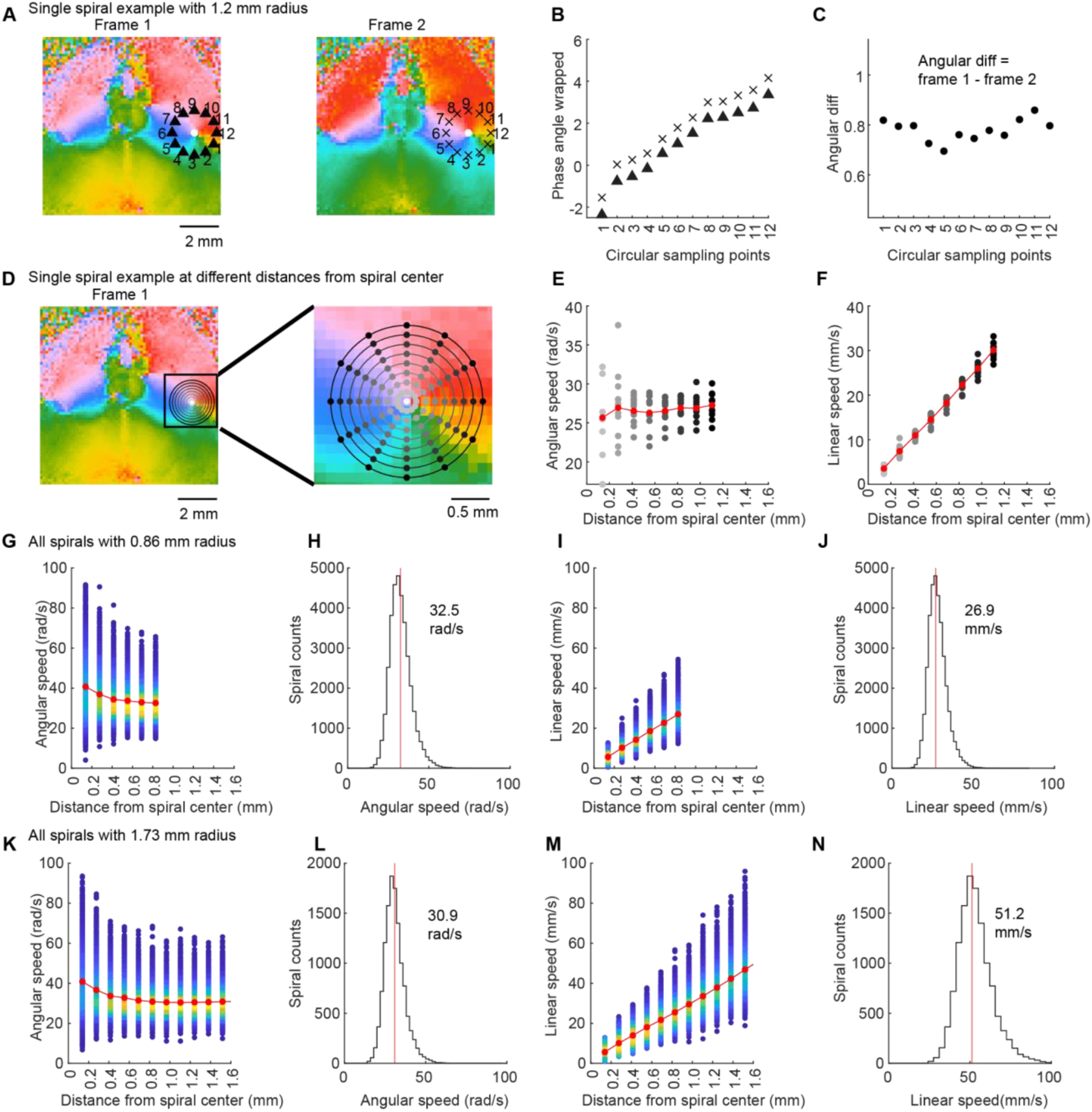
Angular speed and linear speed calculation for spirals. To avoid the estimation errors of speed from optimization methods such as optical flow, we took advantage of the fact that spiral center coordinates and radius were automatically detected through detection pipeline (see Methods). Angular speed (ω) for a given pixel within the spiral is the angular displacement between 2 nearby frames, while linear speed (v) equals to radius (r) * angular speed (ω). (**A**) Left: An example frame with spiral center and 12 evenly sampled points at spiral maximum radius (1.2mm) overlaid on top of the spiral frame. Right: Next frame after frame on the left. (**B**) Phase angles from the 12 evenly sampled points in frames in A were unwrapped to eliminate phase discontinuities. (**C**) Angular differences /displacements between the 2 example frames were consistent across all 12 sampling points. Angular speed (rad/s) is the product of angular difference between 2 frames (rad/frame) and frame rate (35 fps). (**D**) Left: Sampling points with incremental distances from the spiral center, up to the maximum radius detected. Right: Zoom in on the spiral center. (**E**) Angular speed (rad/s) for 12 evenly sampled points at incremental distances from the spiral center, up to the maximum radius detected. Different shades of gray color represent different distances from the spiral center as shown in D. Mean angular speed at different distances from the spiral center are overlaid in red. (**F**) Linear velocities calculated from angular velocities in E. Mean linear velocities at different distances from the spiral center are overlaid in red. (**G**) Angular speed density plot for all spirals with 0.86 mm maximum radius detected in all 15 mice, within a 1.7×3.4mm square close to the center of SSp in the right hemisphere. Mean angular speed of all spirals at different distances from the spiral center are overlaid in red. The slight increase in angular speed at a small distance from the spiral center may reflect an increased contribution of noise to these points. (**H**) Angular speed histogram for all spirals with 0.86 mm maximum radius at maximum radius distance. (**I**) Linear speed for all spirals with 0.86 mm maximum radius detected in all 15 animals. Mean linear speed of all spirals at different distances from the spiral center are overlaid in red. (**J**) Histogram of linear speed for all spirals with 0.86 mm radius at maximum radius distance. (**K**) to (**N**) Same as G to J, but for all spirals with 1.73 mm maximum radius detected in all 15 mice.

**Fig. S10.**
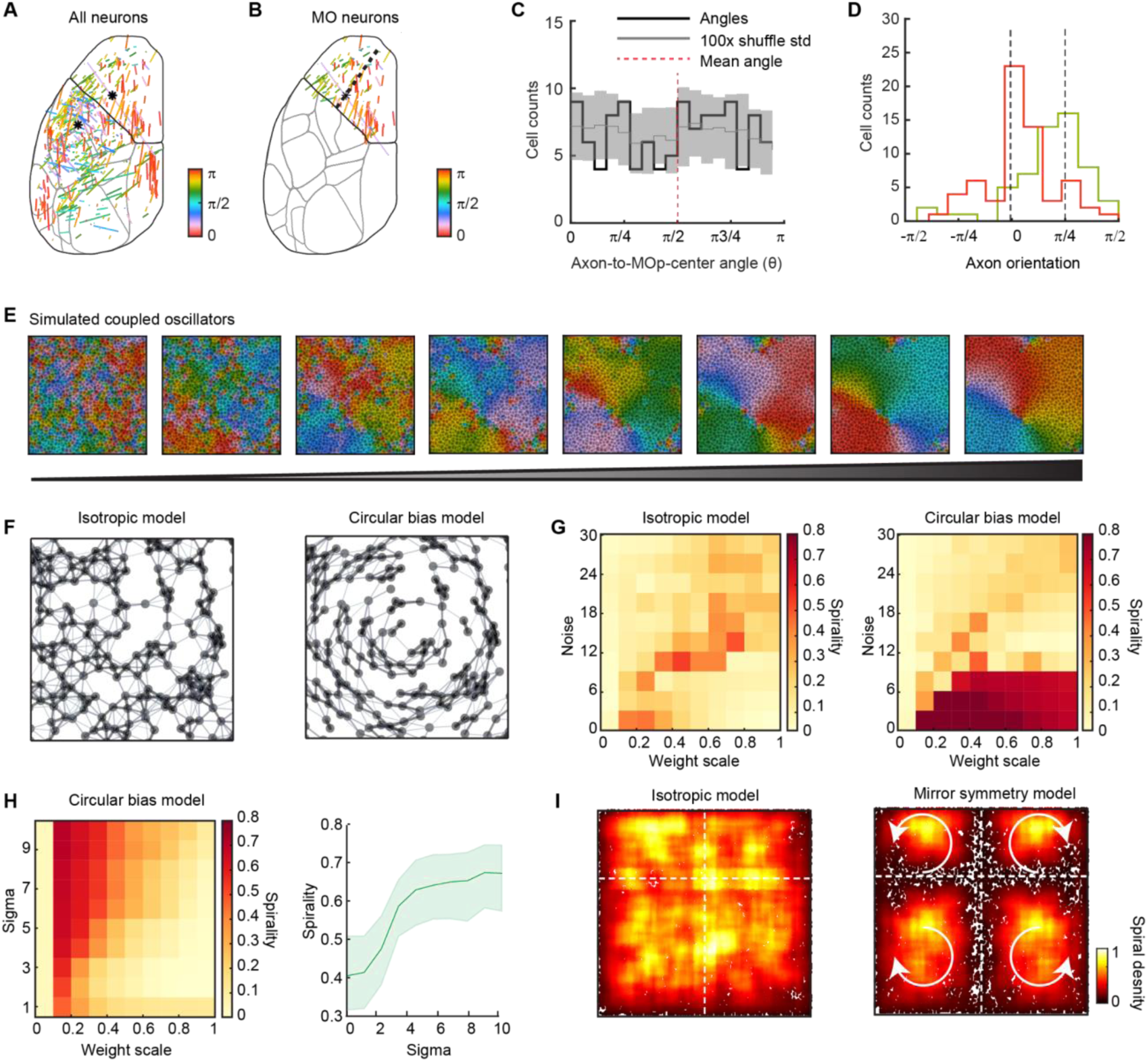
Analysis of axonal architecture in motor cortex, and a coupled oscillator model of spiral dynamics with circular connectivity bias and mirrored topographic connectivity. (**A**) Axonal orientation of all cells within the left hemisphere. (**B**) Axonal orientation of all 120 cells within the motor cortex. (**C**) Axonal orientation of cells within the motor cortex did not form part of a circle, as axon-to-MOp-center angels did not have a distribution peak at 90°, as expected from a circle. This result does not rule out the possibility of a spiral organization in motor neurons, but may require denser morphological sampling. Real data vs permutation distribution: *p* = 0.55, Watson’s U^2^ test. (**D**) Cells below the black dashed line in B had peak biased axonal orientation at 0 (in red), while cells above the black dashed line in B had peak biased axonal orientation at pi/4 (in lime green). Note that axon orientations of all cells are rotated by π/2 in D, compared to B, to avoid distribution discontinuity for visualization purposes. (**E**) An example simulation exhibiting evolution of phase values from a randomized initial condition to spiral pattern over time, following the Kuramoto model. Node positions are plotted with a Voronoi graph. Consistent with previous results (*91*), traveling waves naturally emerged within locally connected networks with exponentially delaying connectivity (Methods). (**F**) Illustration of the connectivity in an isotropic model and circular bias model (Methods). In an isotropic model, a network of coupled oscillators with local connectivity, but no circular bias, was generated to simulate network oscillations. In a circular bias model, enhanced circular connectivity around a centerpoint was imposed. (**G**) Spirality index as a function of weight scale (effective coupling strength) and noise in the isotropic model (left) and circular bias model (right). Spirals emerged within intermediate values of weight scale and noise level in the isotropic model. Spiral waves were characterized with a spirality index (0 no spiral, 1 spiral-like) for spatially coupled oscillators (Methods). Spiral waves were more likely to be observed near the transition between a desynchronized regime and a synchronized regime, consistent with stable vortices/defects during BKT phase transition (*93*, *94*). However, spirals were stabilized even within the higher weight scale (0.4-1) in the circular bias model compared to the isotropic model, with the same initial conditions. Sigma value (circular bias in connectivity) was set at 50 in the circular bias model. (**H**) Left: spirality index increased as a function of weight scale and sigma values (circular bias in connectivity), with noise set at 0. Right: spirality index increased as a function of sigma (r = 0.27, *p* = 4.3 x10^-13^, Pearson correlation), with weight scale chosen at 0.1 and 0.2 from the left heat map. (**I**) To explore how mirror symmetry connectivity affects the spiral distribution, we constructed a mirror symmetry model with coupled oscillators (Methods). Spiral density concatenated from 150 simulation seeds with the same initial condition, in isotropic model (left) and mirror symmetry model (right). Both spiral density plots were normalized to the same peak density value. Each dot represents a detected spiral center. An absence of spirals along the borders in the mirror symmetry model was observed, as in widefield imaging data along hemisphere border, as well as the SSp and MO border (Fig. 3). This is likely due to the rapid annihilation of spiral activity along the boundary, as the mirrored symmetry enforces mirrored phase pattern that conflicts with the circular phase distribution required for spiral waves. Activity should instead propagate parallel to this boundary, consistent with the data (fig. S8P).

**Fig. S11.**
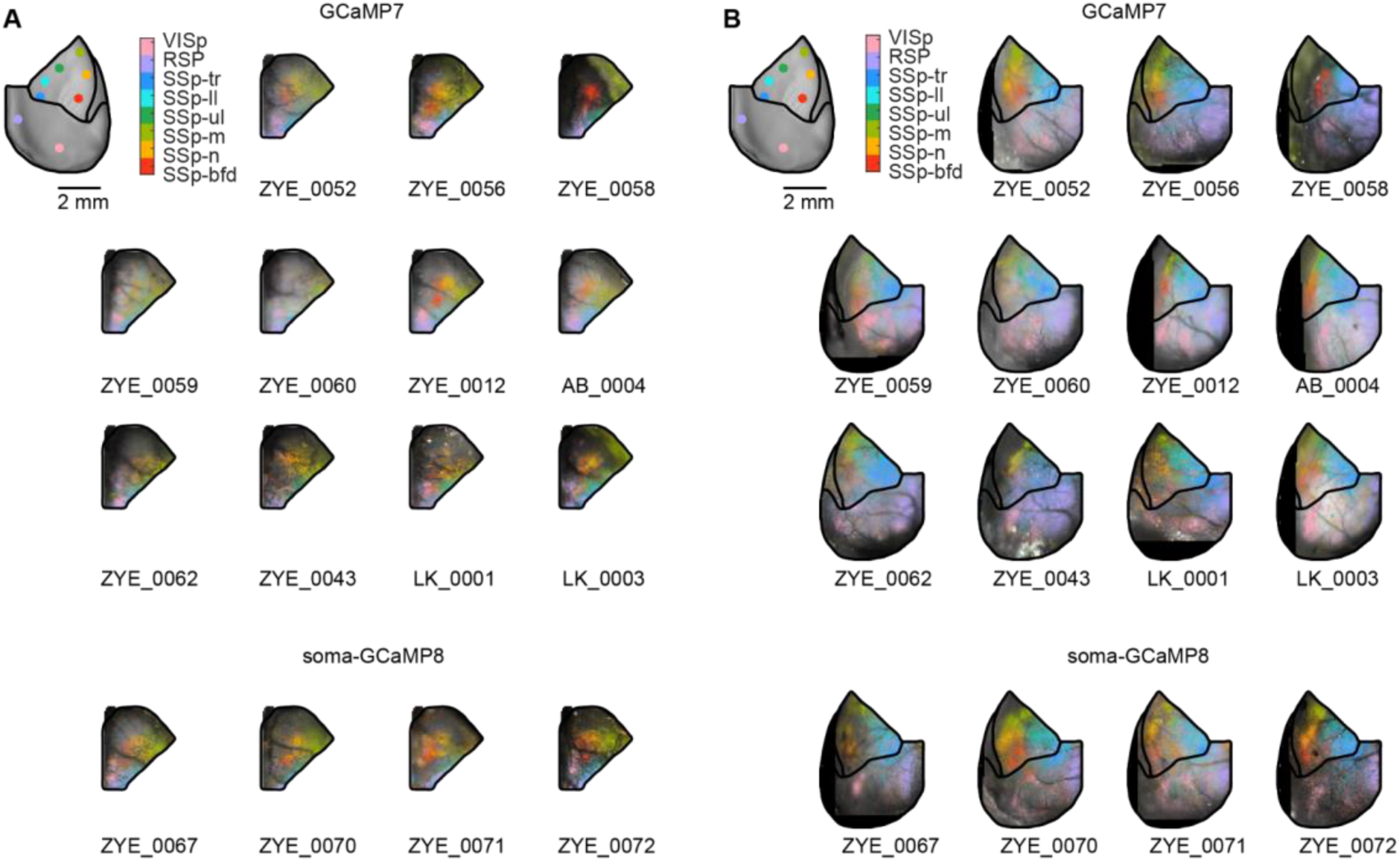
Activity kernel maps were consistent across sessions for motor-to-sensory cortex prediction and left-to-right hemisphere prediction. (**A**) Kernel maps from n=15 mice showing motor-to-sensory cortex predictions for example pixels (filled dots) in the sensory cortex. Top 11 mice: GCaMP7f, bottom 4 mice: GCaMP8s. To account for the fact that fluorescence in GCaMP7s-expressing mice could arise in principle from both somas and axon terminals, we also tested subjects with soma-restricted GCaMP8m expression, which had similar results. (**B**) Same as in A, but for 15 kernel maps from left-to-right hemisphere predictions.

**Fig. S12.**
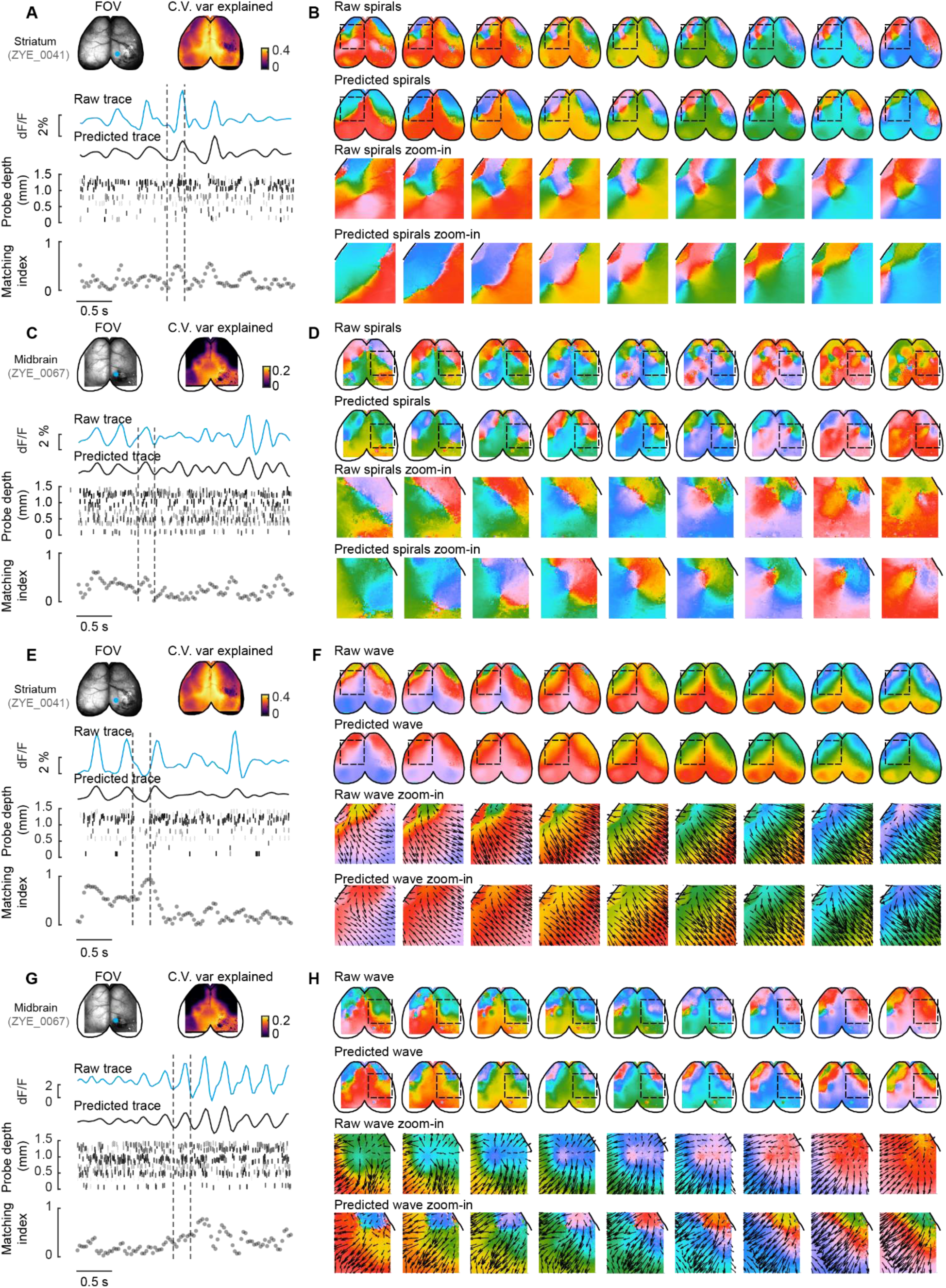
Example spiral and traveling wave predictions from subcortical spiking activity. (**A**) Example epoch of cortical spiral prediction from striatal spiking activity. Top row left: Mean intensity frame of widefield field of view (FOV). Note that the penetration position of the 4-shank Nueropixels probe was from the posterior cortex in the right hemisphere. Top row right: Cross-validated variance explained map of the cortical data from prediction with striatal spiking activity. Second row: Example raw (blue) and predicted (black) cortical activity trace in the example pixel, as indicated in the widefield FOV on the top row (blue dot). Third row: Raster plot of the spiking activity recorded from the Neuropixels probe. Spikes from all 4 shanks were concatenated, and ordered from bottom to top by the depth of the probe from the tip (0 mm) to the surface of the probe correspondingly. Each neuron was color coded by a randomized shade of gray color. Bottom row: Frame-to-frame traveling wave matching index. Note a general increase of wave matching index as 2-8 Hz amplitude increased. (**B**) Example phase maps of the raw (top row) and predicted (second row) cortical activity within the dashed line in A, and the zoom-in of spirals in raw (third row) and predicted (fourth row) phase maps. (**C**), (**D**) Same as in A and B, but for an example midbrain recording session. (**E**) Example epoch of cortical traveling wave prediction from striatal spiking activity. Same session as in A, but in a different time epoch. (**F**) Example phase maps of the raw (top row) and predicted (second row) cortical activity within the dashed line in e, and the zoom-in of the raw (third row) and predicted (fourth row) phase maps and traveling waves, quantified with optical flow (Methods). Note that the directions of the raw and predicted traveling waves were similar, which were quantified as high traveling wave matching index in E. (**G**), (**H**) Same as in E, F, but for an example midbrain recording session. Same session as in c, but in a different time epoch.

**Fig. S13.**
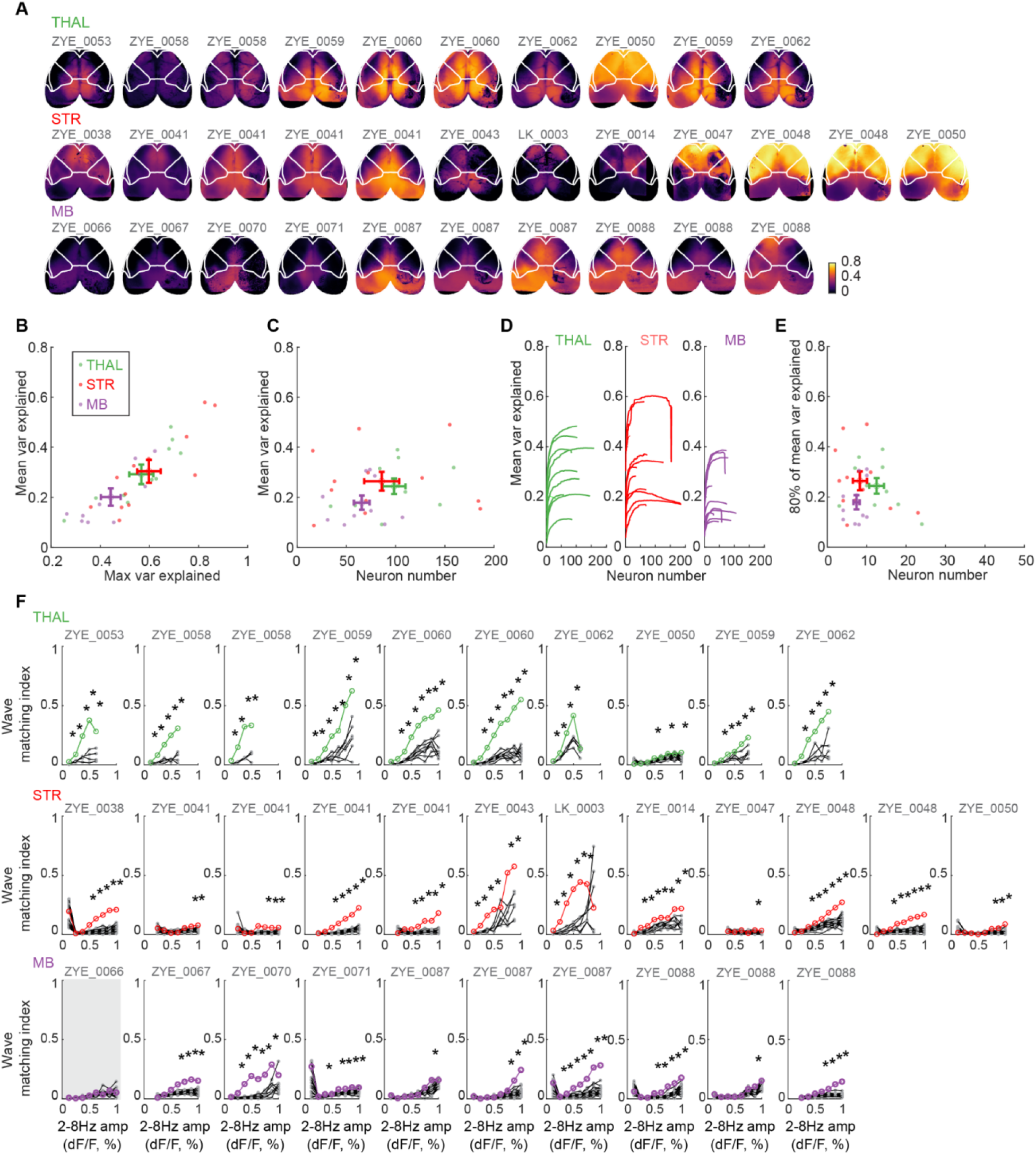
Prediction of cortical widefield activity and traveling waves from subcortical spiking activity. (**A**) Cross-validated variance explained maps for cortical prediction from spiking data in three different subcortical regions. Top: thalamus; middle: striatum; bottom: midbrain. (**B**) Mean variance explained amongst all three regions were not significantly different (thalamus: 29.2 ± 3.9%, 10 sessions; striatum: 30.4 ± 4.6%, 12 sessions; midbrain: 20.1 ± 3.5%, n = 10 session; mean ± SEM; thalamus vs striatum, p = 0.85; thalamus vs midbrain, p = 0.1; striatum vs midbrain, p = 0.1; Student’s t-test, p<0.05). Peak variance explained for midbrain were slightly smaller than thalamus and striatum (thalamus: 56.7 ± 4.9%; striatum: 59.8 ± 4.8%; midbrain: 44.3 ± 4.0%; mean ± SEM; thalamus vs striatum, p = 0.66; thalamus vs midbrain, p = 0.06; striatum vs midbrain, p = 0.02; Student’s t-test, p<0.05). (**C**) Total number of neurons recorded in midbrain is smaller than thalamus (thalamus: 99 ± 12 neurons; striatum: 86 ± 18 neurons; midbrain: 66 ± 8 neurons; mean ± SEM; thalamus vs striatum, p = 0.58; thalamus vs midbrain, p = 0.03; striatum vs midbrain, p = 0.34; Student’s t-test, p<0.05). (**D**) Cumulative variance explained for cortical activity prediction from spiking data in three subcortical regions, ordered by contributions of each single neuron. Left: thalamus; middle: striatum; right: midbrain. In the first round of iteration for each session, cortical widefield activity was first predicted from each single neuron separately, and neurons were then sorted in descending order based on each neuron’s contribution to variance explained. In the second round of prediction iteration for the same session, numbers of neurons used for prediction cumulatively increased from 1 to maximum number of neurons based on the contribution order established earlier, and cross-validated variance explained were calculated for each increment. Note that, in some sessions, variance explained starts to decrease after plateau, as neurons in the bottom contribution ranks start to add noise to full prediction with overfitting. (**E**) Numbers of neurons accounting for 80% of peak cross-validated variance explained in D were similar for all three subcortical regions with slightly higher numbers in thalamus (thalamus: 13 ± 2 neurons; striatum: 8 ± 2 neurons; midbrain: 7 ± 1 neurons; mean ± SEM; thalamus vs striatum, p = 0.12; thalamus vs midbrain, p = 0.02; striatum vs midbrain, p = 0.69; Student’s t-test, p<0.05). (**F**) Wave matching indexes increased as 2-8 Hz amplitude increased, and were significantly above permutation control (Thalamus: 10/10 session; Striatum: 12/12 session; midbrain: 9/10 sessions; unbalanced Two-Way ANOVA test, p<0.05). Session that failed the significance test is highlighted in gray. As examples above demonstrated (fig. S12), wave matching indexes were high when 2-8Hz amplitudes were high. To quantify this relationship, we binned frames based on the mean 2-8 Hz amplitude in the raw cortical widefield data, and calculated the wave matching index from all frames within the amplitude bins. When original and predicted cortical wave directions, quantified with optical flow, matched well for the majority of pixels in frames from a given 2-8 Hz amplitude bin, wave angular difference values amongst all pixels between raw and predicted optical flow were uniformly distributed with low circular variance (close to 0), resulting in a matching index close to 1; when original and predicted cortical wave directions did not match, angular difference values were spread out with high circular variance (close to 1), resulting in a matching index close to 0. For permutation tests, unit identities were shuffled in the test dataset for 20 times without retraining the model, before optical flow and matching indexes were calculated. If the oscillation phase of different units within the subcortical areas were uniform, or random without extra information, measured matching index distribution should not be significantly different from shuffled permutations.

